# Inhibited KdpFABC resides in an E1 off-cycle state

**DOI:** 10.1101/2022.06.19.496728

**Authors:** Jakob M. Silberberg, Charlott Stock, Lisa Hielkema, Robin A. Corey, Jan Rheinberger, Dorith Wunnicke, Victor R. A. Dubach, Phillip J. Stansfeld, Inga Hänelt, Cristina Paulino

**Affiliations:** Institute of Biochemistry, Biocenter, Goethe University Frankfurt, Max-von-Laue-Straße 9, 60438, Frankfurt/Main, Germany; DANDRITE, Nordic EMBL Partnership for Molecular Medicine, Department of Molecular Biology and Genetics, Aarhus University, Universitetsbyen 81, DK-8000 Aarhus C, Denmark; Department of Structural Biology, Groningen Biomolecular Sciences and Biotechnology Institute, University of Groningen, Nijenborgh 7, 9747 AG, Groningen, The Netherlands; Department of Biochemistry, University of Oxford, South Parks Road, Oxford, OX1 3QU, UK; School of Life Sciences & Department of Chemistry, University of Warwick, Coventry, CV4 7AL, UK

## Abstract

KdpFABC is a high-affinity prokaryotic K^+^ uptake system that forms a functional chimera between a channel-like subunit (KdpA) and a P-type ATPase (KdpB). At high K^+^ levels, KdpFABC needs to be inhibited to prevent excessive K^+^ accumulation to the point of toxicity. This is achieved by a phosphorylation of the serine residue in the TGES_162_ motif in the A domain of the pump subunit KdpB (KdpB_S162-P_). Here, we explore the structural basis of inhibition by KdpB_S162_ phosphorylation by determining the conformational landscape of KdpFABC under inhibiting and non-inhibiting conditions. Under turnover conditions, we identified a new inhibited KdpFABC conformation that we termed E1-P tight, which is not part of the canonical Post-Albers transport cycle of P-type ATPases. It likely represents the biochemically described stalled E1-P state adopted by KdpFABC upon KdpB_S162_ phosphorylation. The E1-P tight state exhibits a compact fold of the three cytoplasmic domains and is likely adopted when the transition from high-energy E1-P states to E2-P states is unsuccessful. This study represents a structural characterization of a biologically relevant off-cycle state in the P-type ATPase family and supports the emerging discussion of P-type ATPase regulation by such conformations.

## Introduction

A steady intracellular K^+^ concentration is vital for bacterial cells. Various export and uptake systems jointly regulate bacterial K^+^ homeostasis when facing rapid changes in the environment (Diskowski et al., 2015; Stautz et al., 2021). KdpFABC is a primary active K^+^ uptake system, which is produced when the extracellular K^+^ supply becomes too limited for uptake by less affine translocation systems like KtrAB, TrkAH or Kup. Due to its high affinity for K^+^ (K_M_ = 2 µM) and its active transport, KdpFABC can pump K^+^ into the cell even at steep outward-directed gradients of up to 10^4^, thereby guaranteeing cell survival (Altendorf et al., 1998; Epstein et al., 1993; Rhoads and Epstein, 1977).

The heterotetrametric KdpFABC complex comprises four subunits, namely the channel-like KdpA, a member of the superfamily of K^+^ transporters (SKT) (Durell et al., 2000), the P-type ATPase KdpB (Hesse et al., 1984), the lipid-like stabilizer KdpF (Gaßel et al., 1999), and KdpC, whose function is still unknown. KdpB consists of a transmembrane domain (TMD) and the characteristic cytoplasmic nucleotide binding (N), phosphorylation (P) and actuator (A) domains. Analogous to all P-type ATPases, KdpB follows a Post-Albers reaction scheme, switching between E1 and E2 states that provide alternating access to the substrate binding site during turnover (Albers, 1967; Huang et al., 2017; Post et al., 1972; Silberberg et al., 2021; Stock et al., 2018; Sweet et al., 2021). Whilst in its outward-open E1 conformation, KdpFABC binds ATP in the N domain and takes up K^+^ ions via the selectivity filter in KdpA, which progress into the intersubunit tunnel connecting KdpA and KdpB. Binding of ATP causes rearrangements of the cytoplasmic domains that result in nucleotide coordination between the N and P domains in the E1·ATP conformation. The γ-phosphate of ATP is coordinated in close proximity to the highly conserved KdpB_D307_ of the P domain (all residue numbers refer to *Escherichia coli* KdpFABC). Upon binding of K^+^ to the canonical binding site (CBS) of KdpB, KdpB_D307_ cleaves off the ATP γ-phosphate *via* a nucleophilic attack, leading to the autophosphorylation of KdpFABC and progression to the energetically unfavorable E1-P conformation. ATP cleavage releases the N domain from the P domain, allowing relaxing rearrangements of the cytoplasmic domains that convert KdpFABC to the inward-open E2-P state, in which K^+^ is released from the CBS to the cytoplasm due to conformational changes in KdpB’s TMD. Finally, KdpB_D307_ is dephosphorylated by a water molecule coordinated by KdpBE161 of the TGES_162_ loop in the A domain, recycling KdpFABC via the non-phosphorylated E2 state back to its E1 ground state (Huang et al., 2017; Pedersen et al., 2019; Stock et al., 2018; Sweet et al., 2021). Thus, the relative orientation of the three cytoplasmic domains to each other and their nucleotide state (nucleotide-free, nucleotide-bound, or phosphorylated) are crucial for the assignment of catalytic states (Bublitz et al., 2010; Dyla et al., 2020). Notably, while the general catalytic reaction and conformational arrangements of the N, P and A domains follow the conventional Post-Albers cycle observed for other P-type ATPases, the alternating access of the substrate binding site in the KdpFABC complex is inverted to accommodate KdpFABC’s unique inter-subunit transport mechanism involving KdpA and KdpB (Damnjanovic et al., 2013; Silberberg et al., 2021; Stock et al., 2018).

Being a highly efficient emergency K^+^ uptake system, KdpFABC needs to be tightly regulated in response to changing K^+^ conditions, as both too low and too high potassium concentrations would be toxic (Roe et al., 2000; Stautz et al., 2021). At low K^+^ conditions, transcription of the *kdpFABC* operon is activated by the K^+^-sensing KdpD/KdpE two-component system (Polarek et al., 1992). Further, post-translational stimulation is conferred by cardiolipin, whose concentration increases as a medium-term response to K^+^ limitation (Schniederberend et al., 2010; Silberberg et al., 2021). Once K^+^ stress has abated ([K^+^_external_] > 2 mM), the membrane-embedded KdpFABC is rapidly inhibited to prevent excessive uptake of K^+^ (Roe et al., 2000). This is achieved by a post-translational phosphorylation of KdpB_S162_ (yielding KdpFABS162-PC), which is part of the highly conserved TGES_162_ motif of the A domain (Huang et al., 2017; Sweet et al., 2020). In the crystal structure of KdpFABC [5MRW], phosphorylated KdpB_S162_ forms salt bridges with KdpB_K357_ and KdpB_R363_ of the N domain and adopts an unusual E1 conformation. It was suggested that the salt bridge formation inhibits KdpFABC by locking the complex in this conformation (Huang et al., 2017). However, the salt bridges were shown to be non-essential to the inhibition mechanism, leaving the role of this conformation unclear (Stock et al., 2018; Sweet et al., 2020). Recent functional studies showed that KdpB_S162_ phosphorylation stalls the complex in an intermediate E1-P state, preventing the transition to the inward-open E2-P conformation (Sweet et al., 2020).

Here, we set out to address the structural basis for KdpFABC inhibition by KdpB_S162_ phosphorylation. The conformational landscape of KdpFABC was probed under different conditions by cryo-EM, yielding 10 structures representative of 6 distinct states that describe the full conformational spectrum of the KdpFABC catalytic cycle and resolve the effect of the inhibitory phosphorylation on the conformational plasticity of the complex. Distinct conformational states were further characterized by pulsed EPR measurements and MD simulations to decipher how KdpB_S162_ phosphorylation leads to the inhibition of the complex in the high-energy E1-P intermediate.

## Results

Previous structural studies of KdpFABC were exclusively conducted in the presence of conformation-specific inhibitors (Huang et al., 2017; Silberberg et al., 2021; Stock et al., 2018; Sweet et al., 2021). To describe the structural effects of KdpFABC inhibition by KdpB_S162_ phosphorylation, we prepared cryo-EM samples of KdpFABC variants under conditions aimed at covering the conformational landscape of both the non-phosphorylated, active and the phosphorylated, inhibited complex (Table 1; Table 1 – Table Supplement 1). For this, all KdpFABC variants were produced in *Escherichia coli* at high K^+^ concentration, which is known to lead to the inhibitory KdpB_S162_ phosphorylation (Sweet et al., 2020). Then, KdpFABS162AC, a non-phosphorylatable KdpB_S162_ variant, and wild type KdpFABC (KdpFABS162-PC) were analyzed under turnover conditions, i.e., in the presence of saturating KCl and ATP concentrations, to gain insights into the dynamic conformational landscape adopted by the complex. To supplement these samples, inhibited WT KdpFABS162-PC was prepared in the presence of orthovanadate, known to normally arrest P-type ATPases in an E2 state. Further, the catalytically inactive variant KdpFABS162-P/D307NC was prepared under nucleotide-free conditions to investigate E1 apo states. In sum, the 10 maps obtained from the four samples cover the entire KdpFABC conformational cycle. All structures exhibit the same intersubunit tunnel described previously, which varies in length depending on the state, and is filled with densities assigned as K^+^ ions (Silberberg et al., 2021; Stock et al., 2018). Between structures, the ion densities vary slightly in number and position (Table 1 – Figure Supplement 1).

**Table 1:** Conformational landscape of KdpFABC resolved by cryo-EM. Four samples were prepared for cryo-EM analysis. Structural models were built for all maps with a resolution of 4 Å or below.

| Protein sample | Condition | State/Resolution |  |  |  |  |  |  |
| --- | --- | --- | --- | --- | --- | --- | --- | --- |
|  |  | E1 apo open |  | E1 apo tight | E1·ATP<sub>early</sub> | E1-P·ADP | E1-P tight | E2-P |
|  |  | Sub-state 1 | Sub-state 2 |  |  |  |  |  |
| KdpFAB<sub>S162-P</sub>C | 50 mM KCl<br>2 mM ATP<br>("turnover") |  |  |  | 3.5 Å | 3.1 Å | 3.4 Å |  |
|  | 1 mM KCl<br>2 mM VO<sub>4</sub><sup>3-</sup> |  |  |  |  |  | 3.3 Å | 7.4 Å |
| KdpFAB<sub>S162-P/D307N</sub>C | 50 mM KCl | 3.5 Å | 3.7 Å | 3.4 Å |  |  |  |  |
| KdpFAB<sub>S162A</sub>C | 50 mM KCl<br>2 mM ATP<br>("turnover") |  |  |  |  | 3.7 Å |  | 4.0 Å |

**Table 2:** Details of MD simulations run. All simulations were run in POPE membranes, over 3 x 50 ns. RMSDs are the mean and standard deviations over three repeats.

| State | RMSD (nm) |
| --- | --- |
| E1 apo open 1 (nucleotide-free) | 0.47 ± 0.09 nm |
| E1 apo open 2 (nucleotide-free) | 0.55 ± 0.12 nm |
| E1 apo tight (nucleotide-free) | 0.32 ± 0.08 nm |
| E1·ATP<sub>early</sub> (turnover WT) | 0.35 ± 0.04 nm |
| E1-P·ADP (turnover WT) | 0.51 ± 0.04 nm |
| E1-P·ADP (turnover KdpB<sub>S162A</sub>) | 0.30 ± 0.06 nm |
| E1-P tight (turnover WT) | 0.31 ± 0.04 nm |
| E1-P tight (orthovanadate) | 0.27 ± 0.03 nm |

### Non-inhibited KdpFABC transitions through the Post-Albers cycle under turnover conditions

Previous functional and structural studies have shown that non-phosphorylatable KdpFABS162AC can adopt major states of the Post-Albers cycle (Sweet et al., 2021, 2020). However, the different conformations were captured with the help of various state-specific inhibitors, limiting our understanding of KdpFABC’s full conformational landscape. To determine the predominant states under turnover conditions, non-phosphorylatable KdpFABS162AC was incubated with 2 mM ATP and 50 mM KCl for 5 minutes at RT immediately before plunge freezing and analyzed by cryo-EM. From this dataset, we obtained ‘only’ two structures of KdpFABC (Figure 1; Table 1; Figure 1 – Figure Supplement 1–2; Table 1 – Table Supplement 1).

**Figure 1:**
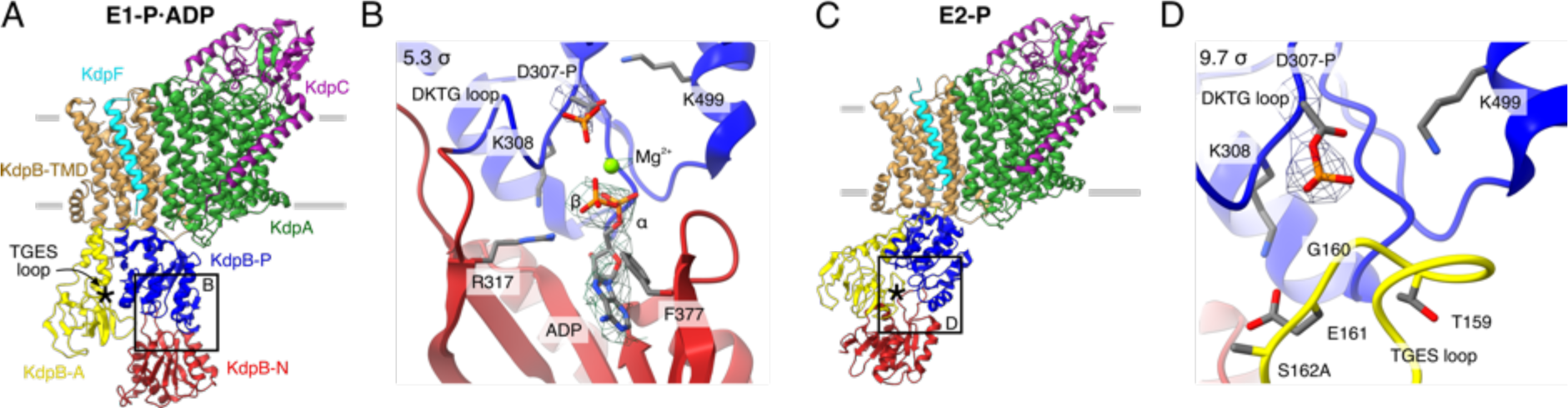
Structures of KdpFABS162AC obtained at turnover conditions. Color code throughout the manuscript, unless stated otherwise, is as follows: KdpF in cyan, KdpA in green, KdpC in purple, KdpB TMD in sand with P domain in blue, N domain in red and A domain in yellow and position of the TGES_162_ loop denoted by an asterisk. **A** E1-P·ADP structure, with its nucleotide binding site (**B)**, showing the bound ADP with Mg^2+^. **C** E2-P structure, with its nucleotide binding site (**D)**, showing the catalytically phosphorylated KdpB_D307_ (P domain), and the TGES_162_ loop (A domain) in close proximity. Densities are shown at the indicated σ level.

The first structure, resolved globally at 3.7 Å and derived from ∼35% of the initial particle set, shows an E1 state. The overall orientation of the cytosolic domains is reminiscent of our previously published E1·ATP structure [7NNL] (Figure 1A; Figure 1 – Figure Supplement 2E) (Silberberg et al., 2021). However, closer inspection of the N domain reveals no density around the expected γ-phosphate of ATP, whereas additional density is observed near KdpB_D307_. We interpret that ADP, instead of ATP, is bound and the catalytic aspartate KdpB_D307_ has been phosphorylated (Figure 1B). An additional density observed near the phosphorylated KdpB_D307_ coincides with an Mg^2+^ ion in the E1-P·ADP SERCA structure [1T5T] (Sørensen et al., 2004), and has thus likewise been assigned as a coordinating Mg^2+^ ion (Figure 1B). Altogether, this conformation represents an E1-P·ADP state, following ATP hydrolysis but preceding the release of ADP. The second structure obtained from the KdpFABS162AC sample under turnover conditions was resolved globally at 4.0 Å from ∼28% of the initial particle set. In this structure, KdpB_D307_ is phosphorylated and the TGES_162_ loop is in close proximity, which is the hallmark of a E2-P conformation (Figure 1C,D). This resembles the previously published E2-P structure [7BH2], which was stabilized by the phosphate analogue BeF3^-^ (Figure 1 – Figure Supplement 2J) (Sweet et al., 2021). In comparison to the E1-P·ADP state, the A domain in the E2-P state has undergone a tilt of 60° and a rotation of 64° around the P domain, positioning the TGES_162_ loop to dephosphorylate KdpB_D307_ in the P domain, while the P domain is also tilted by 40° (Figure 1 – Figure Supplement 3).

The E1-P·ADP and E2-P structures obtained for KdpFABS162AC confirm that, in the absence of the inhibitory KdpB_S162_ phosphorylation, KdpFABC progresses through the entire Post-Albers cycle under turnover conditions, with the ADP release in the E1 conformation and the dephosphorylation of KdpB_D307_ in the E2 conformation likely being the rate-limiting steps that lead to an accumulation of the observed states.

### Inhibited E1-P KdpFABC adopts an off-cycle conformation

To study the structural implications of KdpFABC inhibition by KdpB_S162_ phosphorylation, we next analyzed the conformational landscape of WT KdpFABS162-PC under turnover conditions. To our surprise, the dataset disclosed a higher degree of conformational variability than KdpFABS162AC, yielding three distinct cryo-EM structures (Figure 2; Table 1; Figure 2 – Figure Supplement 1–2; Table 1 – Table Supplement 1). The obtained structures show significant deviations in the cytoplasmic region of KdpB, indicative of different positions in the conformational cycle.

**Figure 2:**
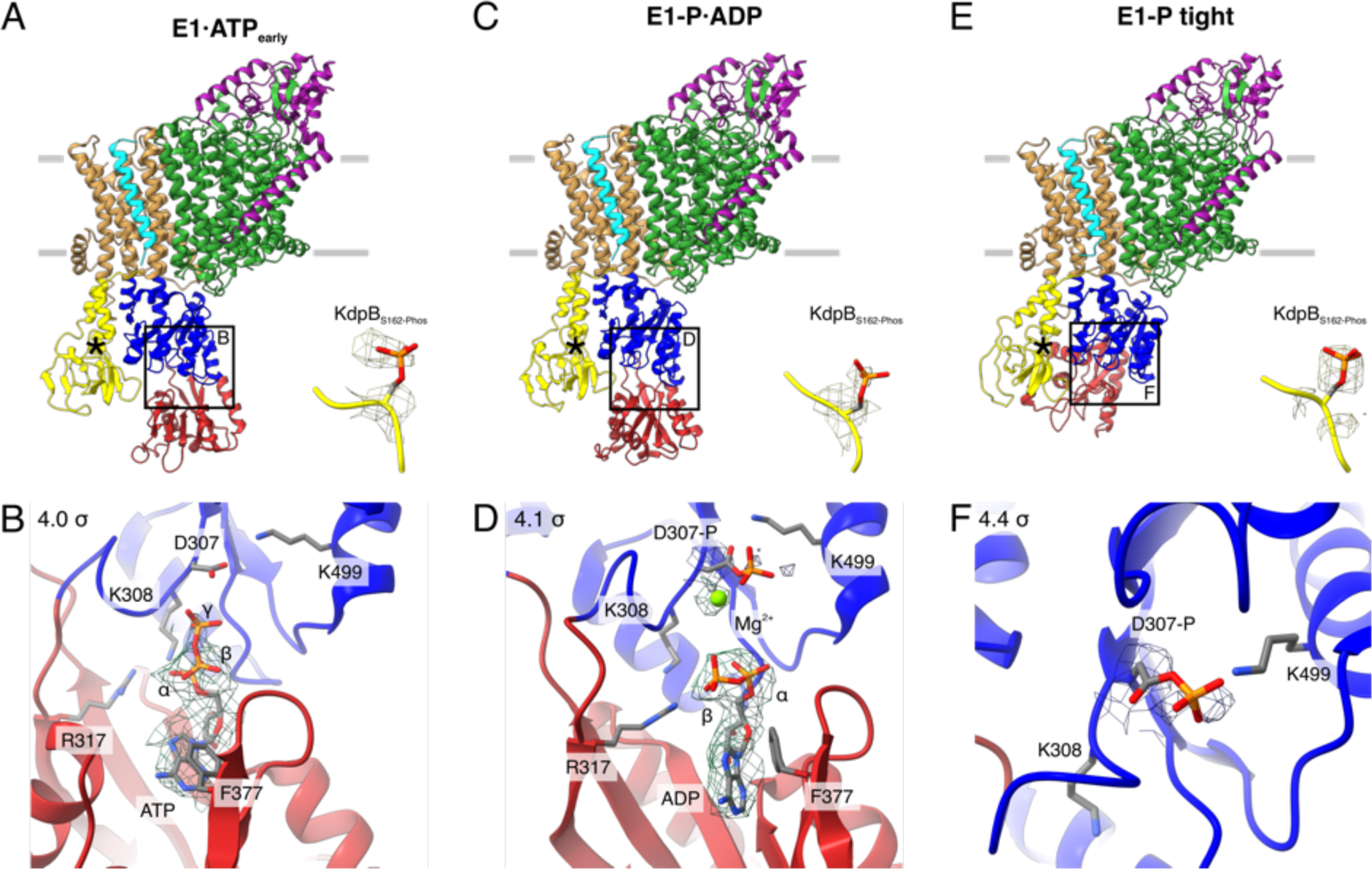
Structures of WT KdpFABS162-PC obtained at turnover conditions. **A** E1·ATP_early_ structure with the corresponding density of KdpB_S162_ phosphorylation and (**B)** its nucleotide binding site, showing the bound ATP. **C** E1-P·ADP structure with the corresponding density of KdpB_S162_ phosphorylation and (**D)** its nucleotide binding site, showing the bound ADP with Mg^2+^. Densities are shown for KdpB_D307-P_, Mg^2+^, and ADP. **E** E1-P tight structure with the corresponding density for KdpB_S162_ phosphorylation and (**F)** its nucleotide binding site. Densities are shown at the indicated σ level.

The first KdpFABS162-PC turnover structure, resolved at an overall resolution of 3.5 Å and derived from ∼9% of the initial particle set, resembles the E1·ATP structure [7NNL] (Silberberg et al., 2021) (Figure 2A; Figure 2 – Figure Supplement 2E). However, a closer analysis of the cytosolic domains and the bound nucleotide reveals significant differences. While ATP is coordinated in a similar fashion as previously observed (Silberberg et al., 2021; Sweet et al., 2021), the N domain is slightly displaced relative to the P domain, providing more access to the nucleotide binding site (Figure 2A,B; Figure 2 – Figure Supplement 2E). We interpret this structure as an E1·ATP conformation at an early stage of nucleotide binding and refer to it as E1·ATP_early_. By contrast, inhibitors such as AMPPCP likely stabilize the latest possible and otherwise transient E1·ATP state right before hydrolysis, explaining the discrepancy between the AMPPCP-stabilized [7NNL] and the E1·ATP state obtained here under turnover conditions. The second KdpFABS162-PC turnover structure, resolved at an overall resolution of 3.1 Å and representing ∼31% of the initial particle set, is virtually identical to the above-described E1-P·ADP conformation of KdpFABS162AC (Figure 2C,D; Figure 2 – Figure Supplement 2J).

While the first two structures represent known states of the catalytic cycle, the third KdpFABS162-PC structure obtained under turnover conditions shows an unusual compact conformation of the cytosolic domains not yet observed in the Post-Albers cycle of other P-type ATPases (Figure 2E,F). The structure was resolved at an overall resolution of 3.4 Å and derived from ∼14% of the initial particle set. Strikingly, the N domain is closely associated with the A domain, thereby disrupting the nucleotide binding site between the N and P domains (Figure 2E,F; Figure 2 – Figure Supplement 3). Closer inspection of the nucleotide binding site shows that the catalytic aspartate KdpB_D307_ is phosphorylated, but not located in proximity to the TGES_162_ loop of the A domain (Figure 2F). This indicates that the conformation is adopted after the canonical E1-P state, which in a normal non-inhibited catalytic cycle would transition into an E2-P state. Due to its compact organization, we termed this conformation E1-P tight.

Previous biochemical studies have shown that KdpB_S162_ phosphorylation inhibits KdpFABC by preventing the transition to an E2 state and stalling it in an E1-P state (Sweet et al., 2020). In line with these observations, we could not identify any conformation following E1-P in the Post-Albers cycle for KdpFABS162-PC, despite the turnover conditions used. Based on this, we hypothesized that the novel E1-P tight state observed represents a non-Post-Albers conformation that is adopted because KdpFABS162-PC cannot proceed to the E2-P state. To further put this to test, we analyzed the conformations adopted by WT KdpFABS162-PC in the presence of the phosphate mimic orthovanadate, which has been shown to trap P-type ATPases in an E2-P conformation and, of all E2 state inhibitors, best mimics the charge distribution of a bound phosphate (Table 1; Figure 2 – Figure Supplement 4–5; Table 1 – Table Supplement 1) (Clausen et al., 2016; Pedersen et al., 2019). Interestingly, KdpFABS162-PC incubated with 2 mM orthovanadate prior to cryo-EM sample preparation did not conform to this behavior. Instead, the major fraction of this sample (∼62% of the initial particle set) resulted in an E1-P tight conformation, resolved at 3.3 Å, which is virtually identical to the E1-P tight conformation obtained under turnover conditions (Figure 2 – Figure Supplement 3H). The position of the orthovanadate coincides with that adopted by the phosphorylated KdpB_D307_, verifying the assignment of this state as a phosphorylated E1-P intermediate (Figure 2 – Figure Supplement 5). Only a minor fraction of the orthovanadate-stabilized sample (∼11% of the initial particle set) adopts an E2-P conformation. Despite the rather poor resolution of 7.4 Å, the conformational state could be confirmed by comparison of the cryo-EM map with the previously published E2-P structure [7BH2] (Figure 2 – Figure Supplement 4I) (Sweet et al., 2021). The minor fraction found in an E2 state likely represents the residual KdpFABC in the sample lacking KdpB_S162_ phosphorylation, as it is in good agreement with the residual ATPase activity level (Sweet et al., 2020). The fact that, in the presence of orthovanadate, KdpFABS162-PC adopts the E1-P tight state instead of an E2-P conformation strongly supports the idea that the inhibited complex cannot transition into an E2 state but instead adopts an off-cycle E1-P state after ADP dissociation.

### A tight conformation is also formed in the absence of nucleotide

Interestingly, the close association of the N domain with the A domain in the E1-P tight conformation also shows some similarities to the E1 crystal structure of KdpFABC [5MRW], the first structure of KdpFABC ever solved (Huang et al., 2017). Unlike our E1-P tight state, the crystal structure is nucleotide-free and does not contain a phosphorylated KdpB_D307_, raising the question how it fits in the conformational landscape of KdpFABC. To further investigate the role of the tight conformation in the conformational cycle of KdpFABC, we prepared a cryo-EM sample of KdpFABS162-P/D307NC under nucleotide-free conditions. This mutant is catalytically inactive and thus restricted to E1 states preceding ATP hydrolysis. In total, we were able to obtain three distinct structures from this preparation, which we assigned to two states relevant to the transport cycle (Figure 3, Table 1; Figure 3 – Figure Supplement 1–2; Table 1 – Table Supplement 1).

**Figure 3:**
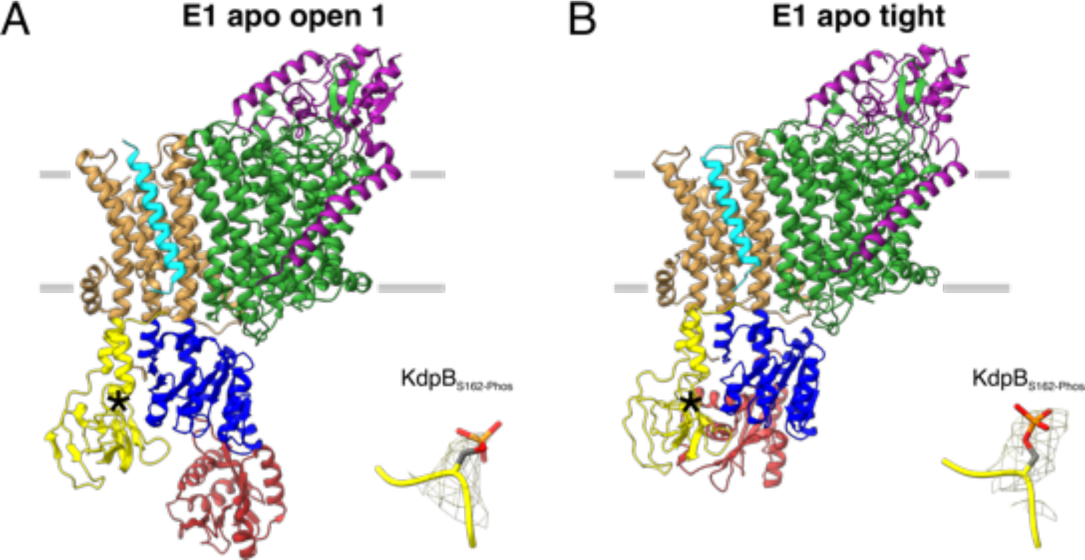
E1 apo structures of KdpFABS162-P/D307NC obtained at nucleotide-free conditions. **A** E1 apo open substate 1 structure with the corresponding density of KdpB_S162_ phosphorylation. The E1 apo open substate 2 differs by a slightly displaced orientation of the N domain (see also Figure 3 – Figure Supplement 1,2). **B** E1 apo tight structure with the corresponding density of KdpB_S162_ phosphorylation.

The first state, composed of 26% of the initial particle set from the nucleotide-free sample, corresponds to the canonical Post-Albers E1 state that precedes ATP binding and resembles the E1 apo state [7BH1] reported previously (Sweet et al., 2021) (Figure 3A; Figure 3 – Figure Supplement 2E). We have termed this conformation the E1 apo open state, as the N and P domains are far apart to provide access to the nucleotide binding site. The cytosolic domains of KdpB reveal a high conformational heterogeneity, evidenced by their lower local resolution (Figure 3 – Figure Supplement 2B,G). Focused 3D classification allowed the distinction of two substates, resolved globally at 3.5 Å and 3.7 Å, differing in the relative position of the N to the P domain (Figure 3 – Figure Supplement 2J). The high degree of flexibility of the N domain in this open conformation likely facilitates nucleotide binding at the start of the Post-Albers cycle. The second state, featuring one structure resolved globally at 3.4 Å and represented by 19% of the initial particle set, shows a compact conformation of the cytosolic domains with the N and A domains in close contact, providing no space for a nucleotide to bind (Figure 3C). This state is similar, but not identical to the one observed in the E1-P tight state and [5MRW], and we have termed it the E1 apo tight state (Figure 3 – Figure Supplement 2O). The presence of a similar tight conformation in both the E1-P and the E1 apo states of KdpB_S162-P_, shows that, while the inhibitory KdpB_S162_ phosphorylation appears to be a prerequisite, the observed close association of the N and A domains can occur before or after the binding and hydrolysis of ATP.

### The tight state interaction between the N and A domains is itself not the cause of inhibition

Structures with a tight arrangement of the A and N domains were only obtained in cryo-EM samples featuring KdpB_S162-P_. To evaluate the dependency of a tight state formation on KdpB_S162_ phosphorylation, we assessed the full conformational flexibility of the N domain using pulsed EPR spectroscopy. Distances between the N and P domains were measured with the labeled residues KdpB_A407CR1_ and KdpB_A494CR1_ (Figure 4A,B; Figure 4 – Figure Supplement 1). Variants were produced at high K^+^ concentrations to confer KdpB_S162_ phosphorylation. EPR analysis of KdpFABS162-P/D307NC in the absence of nucleotide resulted in distances that resemble the E1 apo tight and the E1 apo open conformations (Figure 4B). This corroborates the cryo-EM results that both tight and open states are adopted in the same sample at nucleotide-free conditions. In contrast, the non-phosphorylatable variant KdpB_S162_A resulted in a distance distribution showing a near total elimination of the E1 apo tight state. This strongly indicates that the tight state is stabilized by KdpB_S162_ phosphorylation, although it still exists to a small extent even in the absence of the inhibitory phosphorylation.

**Figure 4:**
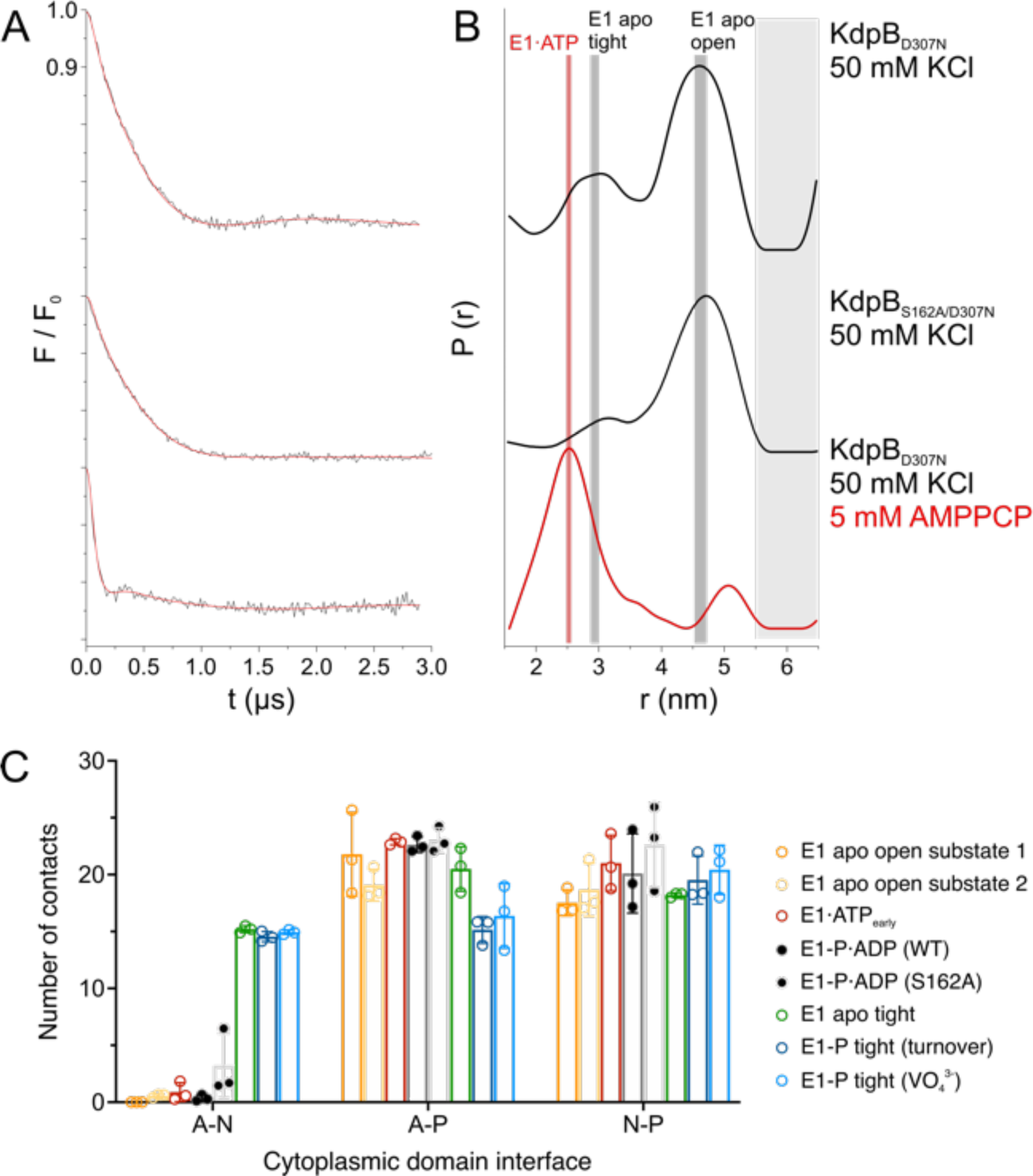
KdpB_S162-P_ increases interdomain contacts in E1 tight states. **A** Background-corrected dipolar evolution function F(t) with applied fit (red lines) of DEER measurements. **B** Area-normalized interspin distance distribution P(r) obtained by Tikhonov regularization. Two KdpFABC variants were prepared with 50 mM KCl and in the presence (red curve) or absence (black curves) of 5 mM AMPPCP. KdpFAB_D307N_C and KdpFAB_S162A/D307N_C without AMPPCP show two conformations, with N-P domain distances of 3 nm and 4.5 nm corresponding to the E1 apo tight and E1 apo open states (dominant state), respectively, as indicated by dark grey background shading. The removal of the inhibitory phosphorylation site in KdpFAB_S162A/D307N_C showed a significant decrease of the distance corresponding to the E1 apo tight state (3 nm) compared to KdpFAB_D307N_C. Addition of AMPPCP to KdpFAB_D307N_C resulted in a single stabilized distance of 2.5 nm, representing the E1·ATP conformation as indicated by red background shading. Light grey shaded areas starting at 5.5 nm indicate unreliable distances. **C** Contact analysis between the N, P and A domains for all E1 structures obtained in this study. Average of the number of contacts (>90% contact) between the different domains over 3 x 50 ns MD simulations (see also Figure 4 – Figure Supplement 2 for details of identified high-contact interactions). Interactions between A and P or N and P domains remain consistent across all states, while interactions between A and N domains are increased only in E1 tight conformations.

To inspect what makes the off-cycle tight conformations energetically preferable, we quantified the cytosolic domain interactions of all E1 states obtained in this study using contact analysis by molecular dynamics (MD) simulation (Figure 4C; Figure 4 – Figure Supplement 2). While no large differences are observed for the contacts between A and P or N and P domains, the off-cycle E1 tight states (E1 apo tight, E1-P tight) are the only ones to show a higher number of interactions between the A and N domains. In fact, they share a nearly identical number of contacts, indicating that the A and N domains move in a concerted manner, similar to previous observations by MD simulations (Dubey et al., 2021). The most evident interaction seen in all three tight states is formed by the salt bridges between the phosphate of KdpB_S162-P_ in the A domain and KdpB_K357/R363_ in the N domain, which were first described in [5MRW] (Huang et al., 2017). However, functional studies have shown that these salt bridges are not essential for the inhibition (Sweet et al., 2020), and neither are they sufficient to fully arrest a tight state, as shown here. In agreement to this, EPR measurements with KdpFAB_S162-P/D307N_C in the presence of the ATP analogue AMPPCP result in a distance distribution that shows a single and well-defined distance, which corresponds to the E1·ATP state. Hence, the salt bridges and the enhanced interaction platform between the A and N domain seen in the tight states likely have a stabilizing role, but can be easily broken and are themselves not the main cause of inhibition.

### E1-P tight is the consequence of an impaired E1-P/E2-P transition

As the compact arrangement found in the E1 tight states itself is not the determining factor of inhibition, the question remains how K^+^ transport is blocked in KdpFAB_S162-P_C and what role the E1-P tight state plays. In a non-inhibited transport cycle, KdpFABC rapidly relaxes from the high-energy E1-P state into the E2-P conformation. To identify what structural determinants enable the arrest before this transition, we compared the E1-P tight structure with the other structures obtained in this study (Figure 5). The E1-P tight state features a tilt of the A domain by 26° in helix 4 (KdpB_198-208_) that is not found in the other E1 states, including the E1 apo tight state (Figure 5B,C – arrow 1). This tilt is reminiscent of the movement the A domain undergoes during the E1-P/E2-P transition (60° tilt, Figure 1 – Figure Supplement 3), but not as far and does not feature the rotation around the P domain (Figure 5 – Figure Supplement 1B,C). Moreover, the P domain does not show the tilt observed in the normal E1-P/E2-P transition. Notably, this is the movement that brings KdpB_D307-P_ in close proximity of KdpB_S162_ in the catalytic cycle (Figure 5 – Figure Supplement 2, Video 1). In an inhibited state, where both sidechains are phosphorylated, such a transition is most likely impaired due to the large charge repulsion. A comparison of the E1-P tight structure with the E1-P·ADP structure, its immediate precursor in the conformational cycle, moreover, reveals a number of significant rearrangements within the P domain (Figure 5B,C). First, Helix 6 (KdpB538-545) is partially unwound and has moved away from helix 5 towards the A domain, alongside the tilting of helix 4 of the A domain (Figure 5B,C – arrow 2). Second, and of particular interest, are the additional local changes that occur in the immediate vicinity of the phosphorylated KdpB_D307_. In the high-energy E1-P·ADP structure, the catalytic aspartyl phosphate, located in the _D307_KTG signature motif, points towards the negatively charged KdpBD518/D522. This repulsion might serve as a driving force for the system to relax into the E2 state in the catalytic cycle. By contrast, the D307KTG loop is largely uncoiled in the E1-P tight state, with the phosphorylated KdpB_D307_ pointing in the opposite direction, releasing this electrostatic strain (Figure 5C – arrow 4). This conformation is further stabilized by a salt bridge formed with KdpB_K499_. The uncoiling in the E1-P tight conformation is likely mediated by the movement of the N domain towards the A domain, as the N domain is directly connected to the _D307_KTG loop (Figure 5C – arrow 3). Altogether, we propose that, in presence of the inhibitory KdpB_S162-P_, the high-energy E1-P·ADP state can no longer transition into an E2-P state after release of ADP and Mg^2+^. As a consequence, the conformational changes observed in the E1-P tight state likely ease the electrostatic tensions of the phosphorylated P domain and stall the system in a ‘relaxed’ off-cycle state.

**Figure 5:**
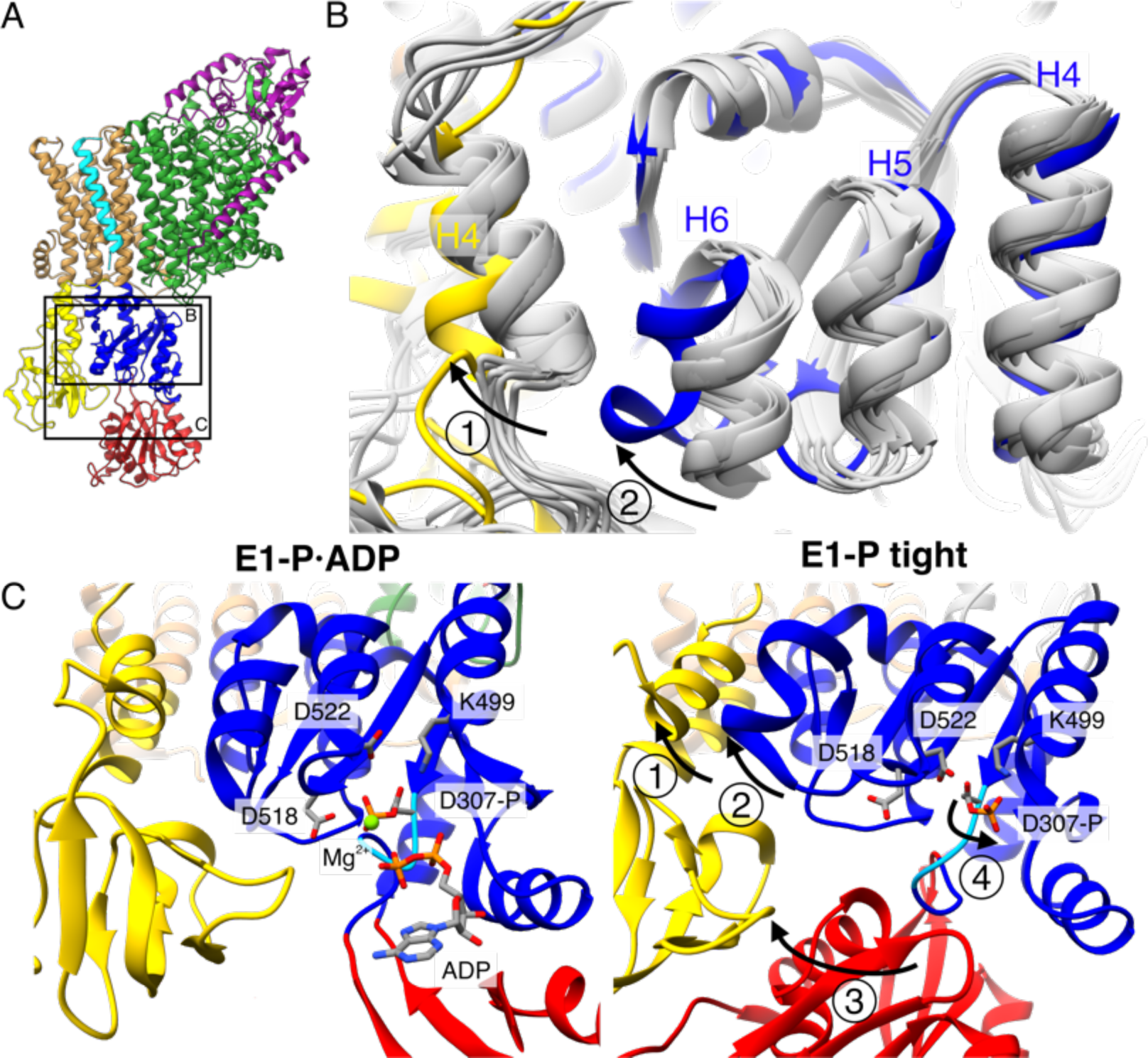
Structural rearrangements in the E1-P tight state facilitate KdpFABS162-PC stalling. **A** Structure of KdpFAB_S162-P_C in the E1-P tight state obtained at turnover conditions. **B** Overlay of all E1 structures determined in this study (in gray) shows helix rearrangements particular to the E1-P tight state (in color). Arrow 1 indicates how the A domain helix 4 tilts away from the TMD by 26°; Arrow 2 indicates how P domain helix 6 moves along with A domain helix 4, slightly uncoiling and shifting away from the remaining P domain. **C** Vicinity of the catalytic aspartyl phosphate (KdpB_D307-P_) in the KdpFAB_S162-P_C E1-P·ADP and E1-P tight structures, showing rearrangements in the P domain to ease tensions of the E1-P state. In the high-energy E1-P·ADP state, KdpB_D307-P_ shows strong electrostatic clashes with KdpB_D518/D522_ (2.9 and 3.5 Å, respectively). These are alleviated by rearrangements in the E1-P tight state. Arrow 3 indicates the movement of the N domain towards the A domain in the tight conformation, which pulls on the _D307_KTG loop (light blue) containing the aspartyl phosphate to uncoil and reorient it. Arrow 4 indicates how this rearrangement reorients the phosphorylated KdpB_D307-P_ away from KdpB_D518/D522_ (7.8 and 4.1 Å, respectively) to form a salt bridge with KdpB_K499_ (3.9 Å).

## Discussion

We set out to deepen our understanding of the structural basis of KdpFABC inhibition via KdpB_S162_ phosphorylation by sampling the conformational landscape under various conditions by cryo-EM. The 10 cryo-EM maps of KdpFABC, which represent six distinct states, cover the full conformational landscape of KdpFABC and, most importantly, uncover a non-Post-Albers regulatory off-cycle state involved in KdpFABC inhibition (Figure 6).

**Figure 6:**
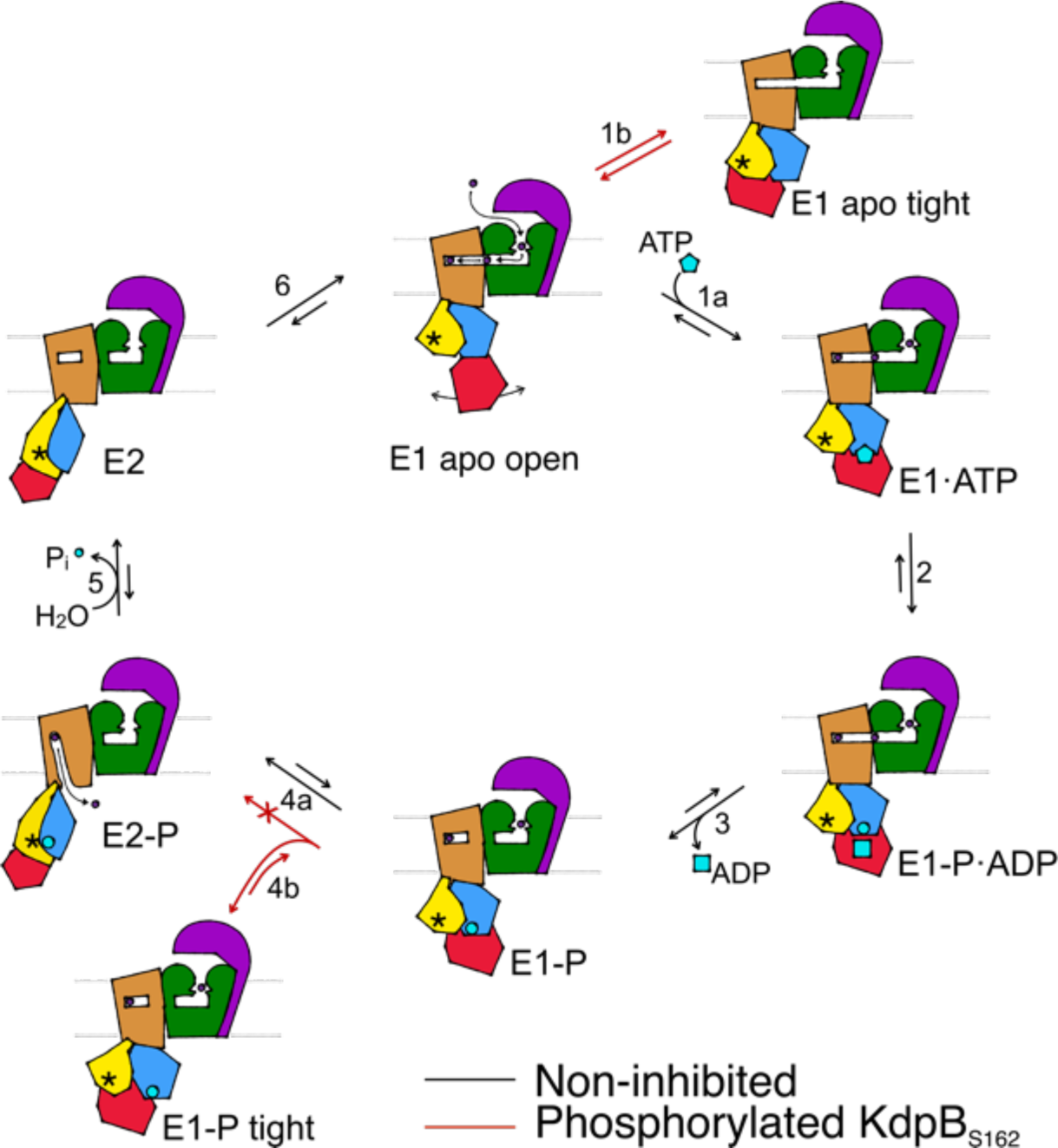
Proposed transport cycle of KdpFABC in the absence and presence of regulatory phosphorylation at KdpB_S162_. Black arrows indicate the non-inhibited catalytic cycle, while red arrows denote the transitions taken when KdpB_S162_ is phosphorylated, and the complex is inhibited. ATP shown as cyan pentagon; ADP as cyan rectangle; phosphorylated KdpB_D307_ as cyan circle; K^+^ ions in dark purple; the position of the TGES_162_ loop in the A domain as an asterisk; KdpF is removed for simplicity.

The catalytic conformations identified here closely adhere to the classical Post-Albers cycle of KdpFABC (Video 1). In the outward-facing E1 apo open state, the nucleotide binding domain is widely accessible, and the N domain shows a high degree of flexibility, which likely enhances the efficiency of nucleotide binding from the environment. Once a nucleotide is bound, the N domain reorients towards the P domain for shared coordination of the nucleotide in the E1·ATP_early_ state (Figure 6, transition 1a). This conformation shows a slightly more open nucleotide-bound state than the previously reported AMPPCP-stabilized E1·ATP state, which likely can progress closer to ATP hydrolysis (Silberberg et al., 2021; Sweet et al., 2021). Subsequently, ATP cleavage leads to the phosphorylation of the catalytic KdpB_D307_ in the P domain, forming the E1-P·ADP intermediate (Figure 6, transition 2), which could be structurally isolated in this study under turnover conditions. The accumulation of the E1-P·ADP state in both turnover samples analyzed in this study suggests that ADP and Mg^2+^ release (Figure 6, transition 3) is the rate-limiting step of KdpFABC turnover. Following the catalytic cycle, the high-energy E1-P state progresses to the E2-P state, whereby large rearrangements bring the conserved TGES_162_ loop in the A domain near the catalytic site of the P domain. This transition is accompanied by conformational changes in the TMD of KdpB, which switches the complex from an outward- to an inward-open state with respect to the CBS (Figure 6, transition 4a). K^+^ is released to the cytosol and KdpB_D307-P_ is dephosphorylated by the TGES_162_ loop (Figure 6, transition 5). Subsequently, the complex cycles back to the E1 apo open state (Figure 6, transition 6).

Whereas the structures obtained for KdpFAB_S162-PC_ in the E1 apo open, E1·ATP, E1-P·ADP and E2-P conformations align well with corresponding states in the Post-Albers cycle of other P-type ATPases, the nucleotide-free E1 apo tight and E1-P tight states do not. Their overall compact fold is facilitated by KdpB_S162-P_, which increases the contacts of the N and A domains. While the compact fold itself is not the main cause of stalling KdpFABC, it might contribute to stabilizing the inhibited complex limiting the innate flexibility of the N domain. Thus, the E1 apo tight state likely has no major physiological relevance, as in presence of ATP, it would progress through the catalytic cycle up to the point of inhibition.

By contrast, we propose that the E1-P tight state is involved in stalling the KdpFABC complex. We suggest that this state represents the biochemically described E1-P inhibited state (Sweet et al., 2020), and is adopted after ADP release from the high-energy E1-P state, when KdpFABC attempts to relax into the E2-P state (Figure 6, transition 4b). This attempt would explain the partial tilt of the A domain in helix 4, observed only in the E1-P tight structure. As suggested before (Sweet et al., 2020), the full relaxation to the E2-P state is however hindered in the inhibited KdpFAB_S162-P_C by a repulsion between the catalytic phosphate in the P domain (KdpB_D307-P_) and the inhibitory phosphate in the A domain (KdpB_S162-P_), which would come in close proximity during the transition from E1-P to E2-P (Figure 6, transition 4a, Video 1). The adopted stalled E1-P tight state is likely an energetically favored conformation between E1-P and E2-P, easing the high-energy constraints around the phosphorylated KdpB_D307_. In good agreement with this hypothesis, KdpFAB_S162-P_C was stabilized in the E1-P tight conformation even in the presence of the inhibitor orthovanadate, known to otherwise stabilize an E2-P state. Notably, orthovanadate has the same charge distribution as the E1-P tight state with phosphorylated KdpB_D307_. By contrast, KdpFAB_S162-PC_ in the presence of AlF_4_^-^, which has a negative charge less, could previously be stabilized in an E2-P state (Stock et al., 2018), likely because the electrostatic repulsion is lower, making the E1-P/E2-P transition more favorable. For a long time, E1-P states of P-type ATPases have been biochemically characterized as high-energy intermediates. However, the structural determinants for this energetic unfavorability have yet to be described. The conformational changes we observe between the E1-P·ADP state and the relaxed E1-P tight conformation offer a first clue as to the regions involved in destabilizing the E1-P state and triggering the transition to the E2-P conformation. In the E1-P tight state, the electrostatic tensions of the E1-P·ADP state are eased by rearrangements of the catalytic aspartate and the surrounding side chains, likely supported by a pulling of the N domain on the _D307_KTG loop as it associates with the A domain during the E1-P to E1-P tight transition. This could be the main stabilizing role of the tight conformation in KdpB_S162-P_ inhibition. Further, the structural differences in the E1-P tight state also involve a movement of helix 6 in the P domain, which is not in the immediate proximity of the catalytic KdpB_D307_ phosphate. This helix is in close proximity to the A domain and next one of the two connections of the P domain to the TMD, which undergo large conformational changes during the E1/E2 transition. Thus, it may be an important element in signaling the ATPase to initiate the transition to E2-P in the TMD from an outward- to an inward-facing state.

The regulation by inactive conformations outside the Post-Albers cycle has been postulated for other P-type ATPases (Dyla et al., 2020). Observation of translocation by the H^+^ ATPase AHA2 on a single-molecule level revealed that the pump stochastically enters inactive states, from which it can return to its active form spontaneously (Veshaguri et al., 2016). These states are adopted from the E1 conformation, similar to the E1 apo tight state observed under nucleotide-free conditions for KdpFABC. However, no full inhibition in the E1-P state was observed for AHA2, indicating a different mechanism. A crystal structure of the Ca^2+^ pump SERCA was also proposed to represent a non-Post-Albers state (Dyla et al., 2020; Toyoshima et al., 2000). However, the organization of the cytosolic domains differs significantly from the tight conformation seen for KdpB, and a physiological role for this structure remains unclear. Altogether, the structural basis for KdpFABC inhibition by KdpB_S162_ phosphorylation described here, describes in detail the involvement of a non-Post-Albers conformation with a clear regulatory role in P-type ATPase turnover. It extends the conformational landscape and strongly supports the emerging idea of non-catalytic off-cycle conformations with important physiological roles.

## Conclusion

The data presented here illuminates the structural basis for KdpFABC inhibition by KdpB_S162_ phosphorylation. We show that stalled KdpFABC adopts a novel conformation, which is either an intermediate of the E1-P/E2-P transition, or a separate state into which the A and P domains relax when the transition to the E2-P state is hindered. Moreover, the N domain in the inhibited KdpFAB_S162-PC_ associates closely with the A domain in a non-Post-Albers conformation that likely further stabilizes the complex during inhibition. These results prove the involvement of off-cycle states in P-type ATPase regulation, and strongly support the burgeoning discussion of non-Post-Albers states of physiological relevance in the conformational landscapes of ion pumps. Further studies will be required to resolve how the phosphorylation of KdpB_S162_ is mediated, and fully illuminate the destabilization of the E1-P state and subsequent transition to the E2-P state.

## Materials and Methods

### Cloning, protein production and purification

*Escherichia coli kdpFABC* (UniProt IDs: P36937 (KdpF), P03959 (KdpA), P03960 (KdpB), and P03961 (KdpC)) and its cysteine-free variant (provided by J.C. Greie, Osnabrück, Germany) were cloned into FX-cloning vector pBXC3H (pBXC3H was a gift from Raimund Dutzler & Eric Geertsma (Geertsma and Dutzler, 2011) (Addgene plasmid # 47068)) resulting in pBXC3H-KdpFABC and pBXC3H-KdpFABCΔCys, respectively. pBXC3H-KdpFABS162AC and pBXC3H-KdpFABD307NC were created from pBXC3H-KdpFABC by site-directed mutagenesis. Plasmids encoding variants used in EPR experiments include pBXC3H-KdpFABD307N/A407C/A494CCýCys and pBXC3H-KdpFABS162A/D307N/A407C/A494CCΔCys, and were created by site-directed mutagenesis based on pBXC3H-KdpFABCβCys.

KdpFABC and KdpFABC variants for structural analysis and pulsed EPR measurements were produced in *E. coli* LB2003 cells (available from the Hänelt group upon request) transformed with the respective plasmids in 12 L of KML (100 μg/ml ampicillin). Cell growth and harvesting was carried out as described previously (Stock et al., 2018). KdpFABC and KdpFABC variants were purified as previously described for wild-type KdpFABC (Stock et al., 2018).

### Cryo-EM sample preparation

#### Wild type KdpFABC under turnover conditions

Purified wild-type KdpFABC was concentrated to 5 mg/ml and supplemented with 50 mM KCl and 2 mM ATP. The sample was incubated at room temperature for 5 min before grid preparation.

#### Wild type KdpFABC stabilized by orthovanadate

Purified wild-type KdpFABC was concentrated to 4 mg/ml and supplemented with 0.2 mM orthovanadate and 1 mM KCl before grid preparation.

#### KdpFAB_S162AC_ under turnover conditions

Purified KdpFAB_S162AC_, in which the inhibitory phosphorylation in KdpB is prevented by the mutation of the phosphorylated KdpB_S162_ to alanine, was concentrated to 3.4 mg/ml and supplemented with 50 mM KCl and 2 mM ATP. The sample was incubated at room temperature for 5 min before grid preparation.

#### KdpFAB_D307N_C under nucleotide-free conditions

Purified KdpFAB_D307N_C, which is prevented from progressing into an E1-P state by the mutation of the catalytic KdpB_D307_ to an asparagine, was concentrated to 4 mg/ml and supplemented with 50 mM KCl before grid preparation.

### Cryo-EM grid preparation

For wild-type KdpFABC with orthovanadate and nucleotide-free KdpFAB_D307N_C, 2.8 µl of sample were applied to holey-carbon cryo-EM grids (Quantifoil Au R1.2/1.3, 200 mesh), which were previously glow-discharged at 5 mA for 20 s. Grids were blotted for 3 - 5 s in a Vitrobot (Mark IV, Thermo Fisher Scientific) at 20 °C and 100% humidity, and subsequently plunge-frozen in liquid propane/ethane and stored in liquid nitrogen until further use. For the turnover samples of wild-type KdpFABC and KdpFAB_S162AC_, 2.8 µl of sample were applied to holey-carbon cryo-EM grids (Quantifoil Au R1.2/1.3, 300 mesh), which were previously twice glow-discharged at 15 mA for 45 s. Grids were blotted for 2 - 6 s in a Vitrobot (Mark IV, Thermo Fisher Scientific) at 4 °C and 100% humidity, and subsequently plunge-frozen in liquid ethane and stored in liquid nitrogen until further use.

### Cryo-EM data collection

Cryo-EM data were collected on a 200 keV Talos Arctica microscope (Thermo Fisher Scientific) equipped with a post-column energy filter (Gatan) in zero-loss mode, using a 20 eV slit, a 100 μm objective aperture, in an automated fashion provided by EPU software (Thermo Fisher Scientific) or serialEM (Mastronarde, 2005; Schorb et al., 2019) on a K2 summit detector (Gatan) in counting mode. Cryo-EM images were acquired at a pixel size of 1.012 Å (calibrated magnification of 49,407×), a defocus range from -0.5 to -2 μm, an exposure time of 9 sec and a sub-frame exposure time of 150 ms (60 frames), and a total electron exposure on the specimen level of about 52 electrons per Å^2^. Data collection was optimized by restricting the acquisition to regions displaying optimal sample thickness using an in-house written script (Rheinberger et al., 2021) and data quality was monitored on-the-fly using the software FOCUS (Biyani et al., 2017).

### Cryo-EM data processing

For all datasets, the SBGrid (Morin et al., 2013) software package tool was used to manage the software packages.

#### KdpFAB_S162AC_ under turnover conditions

Pre-processing of the acquired data was performed as described above, resulting in the selection of 9,170 out of 11,482 images, which were used for further analysis with the software packages cryoSPARC 3.2.0 (Punjani et al., 2017) and RELION 3.1.1 (Zivanov et al., 2018). First, crYOLO 1.7.6 (Wagner et al., 2019) was used to automatically pick 287,232 particles using a loose threshold. Particle coordinates were imported in RELION 3.1.1 (Zivanov et al., 2018), and the particles were extracted with a box size of 240 pixels. Non-protein classes were removed with a single round of 2D classification in cryoSPARC 3.2.0 (Punjani et al., 2017), resulting in 167,721 particles (initial particle set). These particles were then subjected to ab-initio 3D reconstruction in cryoSPARC 3.2.0 (Punjani et al., 2017), and the best two output classes were used in subsequent jobs in an iterative way in RELION 3.1.1 (Zivanov et al., 2018). The dataset was from here on treated separately, with about 37.7% (63,240 particles) in the E2-P state and about 53.3% (89,378 particles) in the E1-P·ADP state. These particles were imported back into RELION 3.1.1, and subjected to 3D classification and refinement, against references obtained for the E1 tight and E1-P·ADP conformations. This resulted in a dataset of 46,904 particles (∼28% of the initial particle set) for the E2-P conformation, and of 70,068 particles for the E1-P·ADP conformation. Several rounds of CTF refinement (Zivanov et al., 2018) were performed, using per-particle CTF estimation. The dataset for the E1-P·ADP conformation was subjected to a round of focused 3D classification with no image alignment, using a mask on the flexible A and N domains of KdpB (Hiraizumi et al., 2019). This resulted in a cleaned dataset of 58,243 particles (∼35% of the initial particle set) for the E1-P·ADP state. In the last refinement iteration, a mask excluding the micelle was used and the refinement was continued until convergence (focused refinement), yielding a final map for the E2-P state at a resolution of 4.3 Å and 4.0 Å after post-processing and masking, sharpened using an isotropic b-factor of -160 Å^2^. The final map for the E1-P·ADP state had a resolution of 4.0 Å after refinement and 3.7 Å after post-processing and masking, and was sharpened using an isotropic b-factor of -123 Å^2^.

#### Wild type KdpFABC under turnover conditions

A total of 17,938 dose-fractionated cryo-EM images were recorded and subjected to motion-correction and dose-weighting of frames by MotionCor2 (Zheng et al., 2017). The CTF parameters were estimated on the movie frames by ctffind4.1.4 (Rohou and Grigorieff, 2015). Bad images showing contamination, a defocus below -0.5 or above -2.0 μm, or a bad CTF estimation were discarded, resulting in 14,604 images used for further analysis with the software packages cryoSPARC 3.2.0 (Punjani et al., 2017) and RELION 3.1.1 (Zivanov et al., 2018). First, crYOLO 1.7.6 (Wagner et al., 2019) was used to automatically pick 1,128,433 particles using a loose threshold. Particle coordinates were imported in RELION 3.1.1 (Zivanov et al., 2018), and the particles were extracted with a box size of 240 pixels. Non-protein classes were removed with a single round of 2D classification in cryoSPARC 3.2.0 (Punjani et al., 2017), resulting in 828,847 particles (initial particle set). These particles were then subjected to ab-initio 3D reconstruction in cryoSPARC 3.2.0 (Punjani et al., 2017), and the best three output classes were used in subsequent jobs in an iterative way in RELION 3.1.1 (Zivanov et al., 2018). The dataset was from here on treated separately, with about 33.2% (275,026 particles) in the E1-P tight state and about 55.1% (346,303 and 110,601 particles) in the E1 nucleotide bound state. These particles were imported back into RELION 3.1.1, and subjected to 3D classification and refinement, against references obtained for the E1 tight and E1·ATP conformations. Several rounds of CTF refinement (Zivanov et al., 2018) were performed, using per-particle CTF estimation, before subjecting all datasets to a round of focused 3D classification with no image alignment, using a mask on the flexible A and N domains of KdpB (Hiraizumi et al., 2019). This resulted in a cleaned dataset of 114,588 particles (∼14% of the initial particle set) for the E1-P tight state, and 277,912 and 80,798 for the E1 nucleotide bound conformations. The latter two were merged and subjected to several rounds of CTF refinement (Zivanov et al., 2018) using per-particle CTF estimation, before subjecting the dataset to another round of focused 3D classification with no image alignment, using a mask on the flexible A and N domains of KdpB (Hiraizumi et al., 2019). This resulted in two distinct datasets of 257,675 particles (∼31% of the initial particle set) for the E1-P·ADP conformation and of 76,121 particles (∼9% of the initial particle set) for the E1·ATP conformation. In the last refinement iteration, a mask excluding the micelle was used and the refinement was continued until convergence (focused refinement), yielding a final map for the E1-P tight state at a resolution of 3.7 Å and 3.4 Å after post-processing and masking, sharpened using an isotropic b-factor of -134 Å^2^. The final map for the E1-P·ADP state had a resolution of 3.4 Å after refinement and 3.1 Å after post-processing and masking, and was sharpened using an isotropic b-factor of -122 Å^2^. The final map for the E1·ATP state had a resolution of 3.9 Å after refinement and 3.5 Å after post-processing and masking, and was sharpened using an isotropic b-factor of -132 Å^2^. No symmetry was imposed during 3D classification or refinement.

#### Wild type KdpFABC with orthovanadate

Pre-processing of the acquired data was performed as described above, resulting in the selection of 2,014 out of 2,488 images, which were used for further analysis with the software package RELION 3.0.8 (Zivanov et al., 2018). First, crYOLO 1.3.1 (Wagner et al., 2019) was used to automatically pick 164,891particles using a loose threshold. Particle coordinates were imported in RELION 3.0.8 (Zivanov et al., 2018), and the particles were extracted with a box size of 240 pixels. Non-protein classes were removed with 2D classification, resulting in 120,077 particles (initial particle set). For 3D classification and refinement, the map of the previously generated E1 conformation EMD-0257 (Stock et al., 2018) was used as reference for the first round, and the two best output classes were used in subsequent jobs in an iterative way. The dataset was from here on treated separately, with about 70.9% (85,102 particles) in the E1-P tight state and about 11.2% (13,508 particles) in the E2-P state. Sequentially, on the E1-P tight dataset several rounds of CTF refinement, using per-particle CTF estimation, and Bayesian polishing were performed (Zivanov et al., 2018), before subjecting the dataset to a round of focused 3D classification with no image alignment, using a mask on the flexible A and N domains of KdpB (Hiraizumi et al., 2019). This resulted in a cleaned dataset of 74,927 particles (∼62% of the initial particle set) for the E1-P tight state, and was subjected to several rounds of CTF refinement, using per-particle CTF estimation, and Bayesian polishing (Zivanov et al., 2018). In the last refinement iteration, a mask excluding the micelle was used and the refinement was continued until convergence (focused refinement), yielding a final map for the E1-P tight state at a resolution of 3.3 Å, and 3.3 Å after post-processing and masking, sharpened using an isotropic b-factor of -55 Å^2^. The final map for the E2-P state (from ∼11% of the initial particle set) had a resolution of 8.7 Å after refinement and 7.4 Å after post-processing and masking, and was sharpened using an isotropic b-factor of -195 Å^2^. No symmetry was imposed during 3D classification or refinement.

#### KdpFAB_D307N_C under nucleotide-free conditions

Pre-processing of the acquired data was performed as described above, resulting in the selection of 12,864 out of 17,889 images, which were used for further analysis with the software package RELION 3.0.8 (Zivanov et al., 2018). First, crYOLO 1.3.1 (Wagner et al., 2019) was used to automatically pick 728,674 particles using a loose threshold. Particle coordinates were imported in RELION 3.0.8 (Zivanov et al., 2018), and the particles were extracted with a box size of 240 pixels. Non-protein classes were removed in several rounds of 2D classification, resulting in 469,466 particles (initial particle set). Due to the large conformational differences between both states, the full dataset was further cleaned by two independent 3D classifications against references obtained for the E1 tight state or the E1 apo open state. Particles belonging to the best classes of both runs were merged and duplicates subtracted, resulting in 306,942 particles that were subjected to a multi-reference 3D classification with no image alignment. The dataset was from here on treated separately, with about 33.7% (158,353 particles) in the E1 tight state and about 31.4% (147,589 particles) in the open state. Several rounds of CTF refinement (Zivanov et al., 2018) were performed, using per-particle CTF estimation, before subjecting both datasets to a round of focused 3D classification with no image alignment, using a mask on the flexible A and N domains of KdpB (Hiraizumi et al., 2019). This resulted in a cleaned dataset of 88,852 particles (∼19% of the initial particle set) for the E1 tight state, 75,711 particles (∼16% of the initial particle set) for the E1 apo open state 1, and 47,981 particles (∼10% of the initial particle set) for the E1 apo open state 2. In the last refinement iteration, a mask excluding the micelle was used and the refinement was continued until convergence (focused refinement), yielding a final map for the E1 tight state at a resolution of 3.8 Å and 3.4 Å after post-processing and masking, sharpened using an isotropic b-factor of -113 Å^2^. The final map for the E1 apo open state 1 had a resolution of 3.9 Å after refinement and 3.5 Å after post-processing and masking, and was sharpened using an isotropic b-factor of -117 Å^2^. The final map for the E1 apo open state 2 had a resolution of 4.0 Å after refinement and 3.7 Å after post-processing and masking, and was sharpened using an isotropic b-factor of -119 Å^2^. For 3D classification and refinement, the map of the previously generated E1 conformation [EMD-0257] (Stock et al., 2018) was used as reference for the first round, and the best output class was used in subsequent jobs in an iterative way. For all datasets, local resolution estimates were calculated by RELION and no symmetry was imposed during 3D classification or refinement. All resolutions were estimated using the 0.143 cut-off criterion (Rosenthal and Henderson, 2003) with gold-standard Fourier shell correlation (FSC) between two independently refined half maps. During post-processing, the approach of high-resolution noise substitution was used to correct for convolution effects of real-space masking on the FSC curve (Chen et al., 2013).

### Model building and validation

Available KdpFABC structures like E1·ATP conformation [7NNL], E1 [6HRA] and E2 conformation [6HRB] were docked into the obtained cryo-EM maps using UCSF Chimera (Pettersen et al., 2004) and used as initial models. Wherever required, rigid body movements were applied to accommodate for conformational changes, and models were subjected to an iterative process of real space refinement using Phenix.real_space_refinement with secondary structure restraints (Afonine et al., 2018; Liebschner et al., 2019) followed by manual inspection and adjustments in Coot (Emsley and Cowtan, 2004). K^+^ ions, cardiolipin, ATP, ADP, P_i_, and orthovanadate were modelled into the cryo-EM maps in Coot. The final models were refined in real space with Phenix.real_space_refinement with secondary structure restraints (Afonine et al., 2018; Liebschner et al., 2019). For validation of the refinement, FSCs (FSC_sum_) between the refined models and the final maps were determined. To monitor the effects of potential over-fitting, random shifts (up to 0.5 Å) were introduced into the coordinates of the final model, followed by refinement against the first unfiltered half-map. The FSC between this shaken-refined model and the first half-map used during validation refinement is termed FSC_work_, and the FSC against the second half-map, which was not used at any point during refinement, is termed FSC_free_. The marginal gap between the curves describing FSC_work_ and FSC_free_ indicate no over-fitting of the model. The geometries of the atomic models were evaluated by MolProbity (Williams et al., 2018).

### EPR sample purification, preparation, data acquisition and analysis

KdpFAB_A407C/A494C_CΔCys, KdpFAB_D307N/A407C/A494C_CΔCys and KdpFAB_S162A/D307N/A407C/A494C_CΔCys, variants based on an otherwise Cys-less background, were produced and purified as described for KdpFAB_G150C/A407C_CΔCys (Stock et al., 2018). Purified and spin-labeled KdpFABCΔCys variants were concentrated to 4 - 7 mg ml^−1^ and supplemented with 14% deuterated glycerol (v/v) and 50 mM KCl. 5 mM AMPPCP stabilizing the E1-ATP conformations was added when indicated.

Pulsed EPR measurements were performed at Q band (34 GHz) and -223 °C on an ELEXSYS-E580 spectrometer (Bruker). For this, 15 μl of the freshly prepared samples were loaded into EPR quartz tubes with a 1.6 mm outer diameter and shock frozen in liquid nitrogen. During the measurements, the temperature was controlled by the combination of a continuous-flow helium cryostat (Oxford Instruments) and a temperature controller (Oxford Instruments). The four-pulse DEER sequence was applied (Pannier et al., 2011) with observer pulses of 32 ns and a pump pulse of 12 - 14 ns. The frequency separation was set to 70 MHz and the frequency of the pump pulse to the maximum of the nitroxide EPR spectrum. Validation of the distance distributions was performed by means of the validation tool included in DeerAnalysis (Jeschke et al., 2006) and varying the parameters “Background start” and “Background density” in the suggested range by applying fine grid. A prune level of 1.15 was used to exclude poor fits. Furthermore, interspin distance predictions were carried out by using the rotamer library approach included in the MMM software package (Jeschke, 2018; Polyhach et al., 2010). The calculation of the interspin distance predictions is based on the cryo-EM structures of the E1 tight, E1 apo open, and E1·ATP conformations for the comparison with the experimentally determined interspin distance distributions.

### Molecular dynamics simulations

Molecular dynamics (MD) simulations were built using the coordinates of eight states of the complex (see Table 3). To reduce the size of the simulation box, KdpA and KdpC were removed from the system, as these were considered unlikely to impact the dynamics of the N, P, and A domains in the timescales simulated. The systems were described with the CHARMM36m force field (Best et al., 2012; Huang et al., 2016) and built into POPE membranes with TIP3P waters and K^+^ and Cl^-^ to 150 mM, using CHARMM-GUI (Jo et al., 2007; Lee et al., 2016). Where present, KdpB_S162_ was phosphorylated for each system, and for the E1·ATP and E1-P·ADP states, the nucleotide was included based on the structural coordinates. Where present, the K^+^ bound in the CBS was preserved.

**Table 3:** EMDB and PDB accession codes of structure depositions.

| State | EMDB # | PDB # |
| --- | --- | --- |
| E1-ATP<sub>early</sub> (turnover WT) | EMD-14913 | 7ZRG |
| E1-P-ADP (turnover WT) | EMD-14917 | 7ZRK |
| E1-P tight (turnover WT) | EMD-14912 | 7ZRE |
| E1-P tight (orthovanadate) | EMD-14911 | 7ZRD |
| E2-P (orthovanadate) | EMD-14347 | N/A |
| E1-P-ADP (turnover KdpB<sub>S162A</sub>) | EMD-14919 | 7ZRM |
| E2-P (turnover KdpB<sub>S162A</sub>) | EMD-14918 | 7ZRL |
| E1 apo tight (nucleotide-free KdpB<sub>D307N</sub>) | EMD-14914 | 7ZRH |
| E1 apo open 1 (nucleotide-free KdpB<sub>D307N</sub>) | EMD-14915 | 7ZRI |
| E1 apo open 2 (nucleotide-free KdpB<sub>D307N</sub>) | EMD-14916 | 7ZRJ |

**Table 4:**
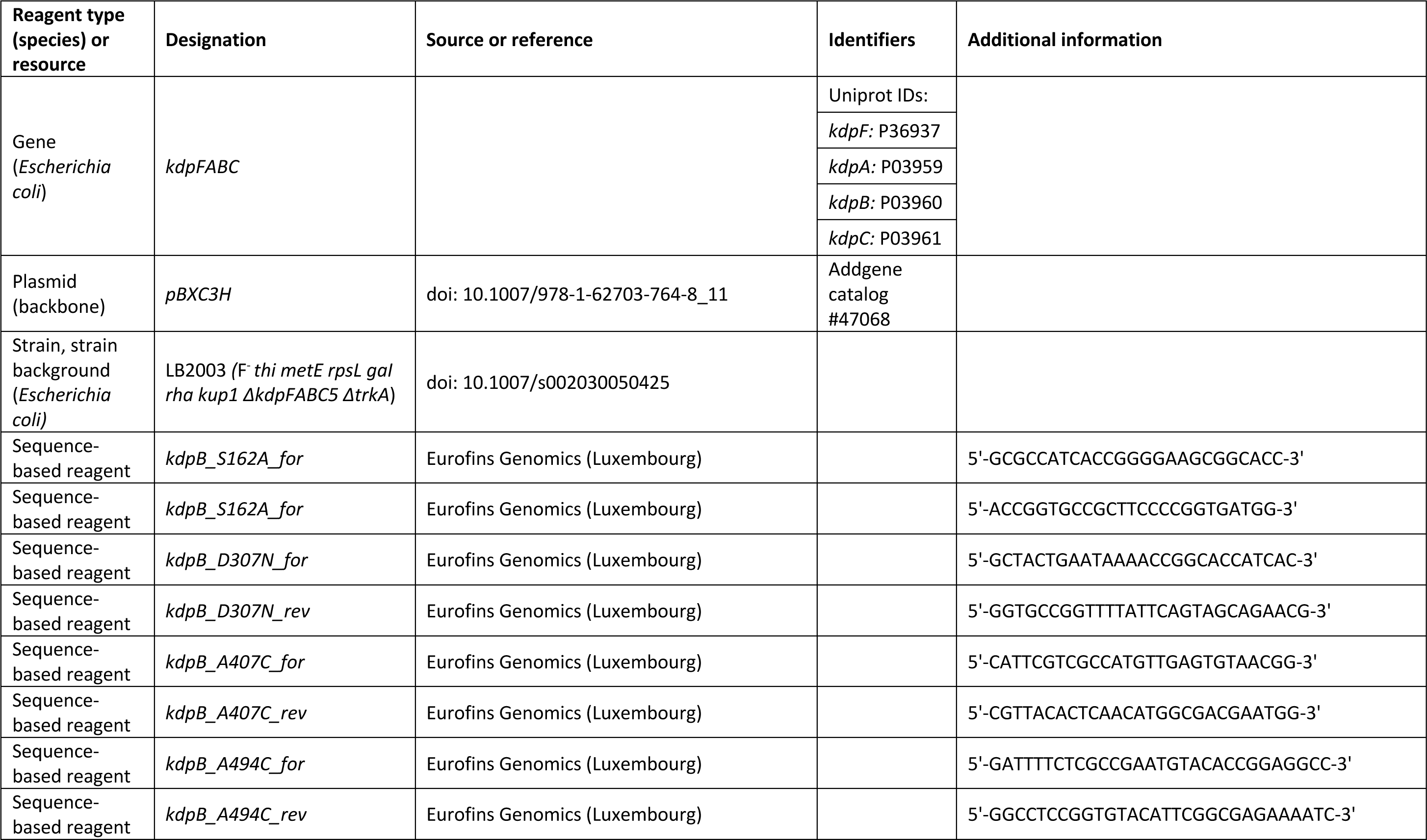
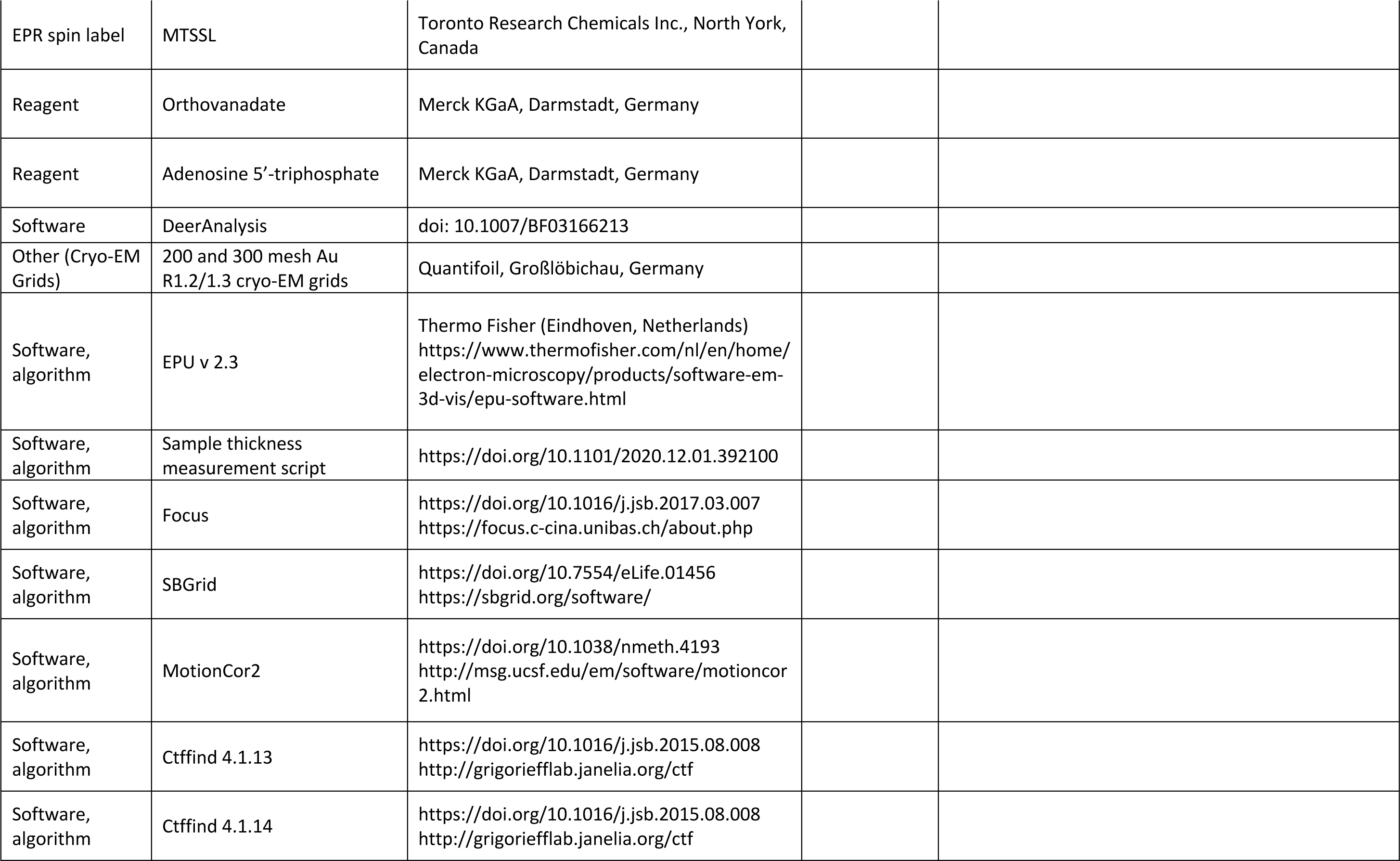
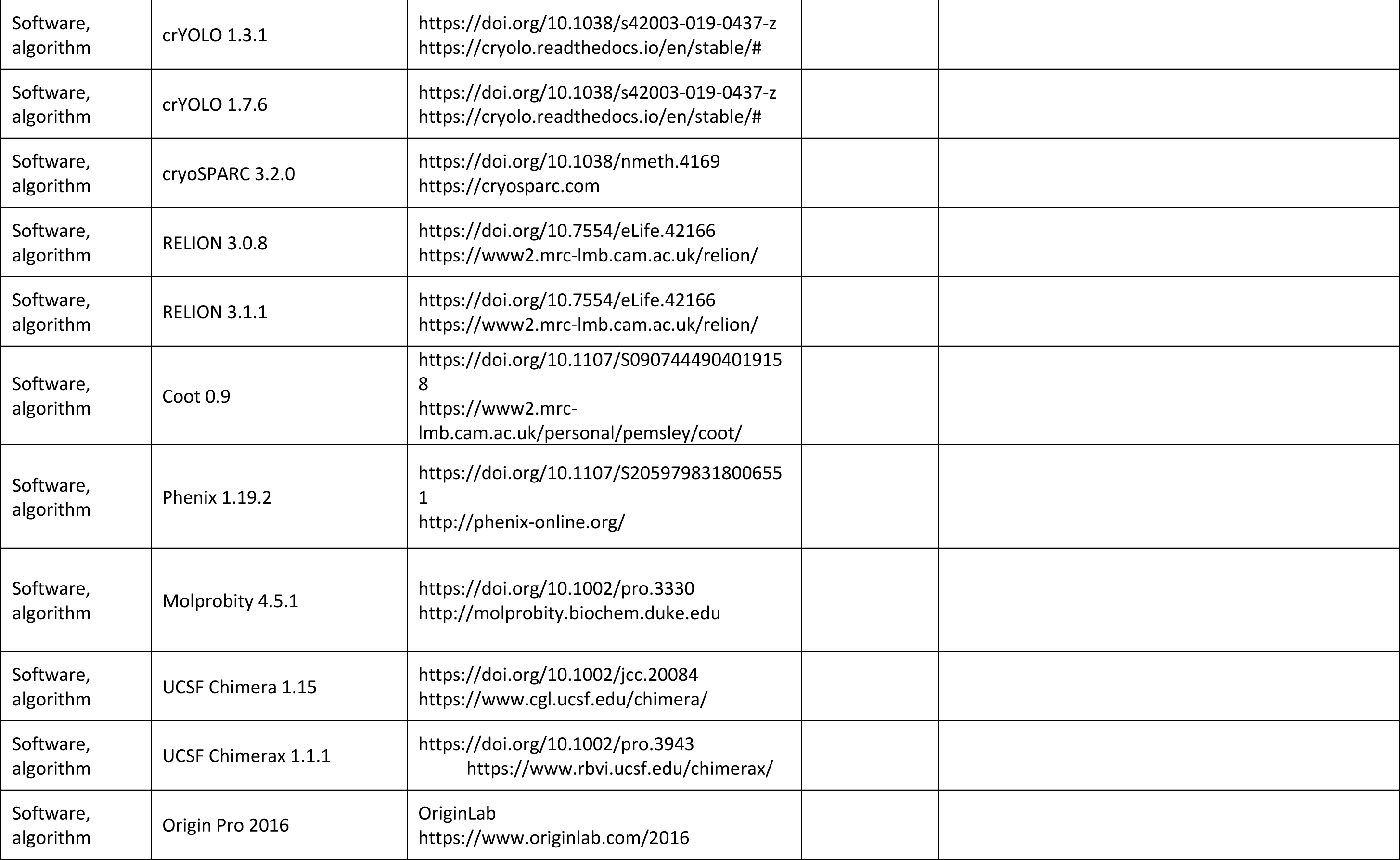
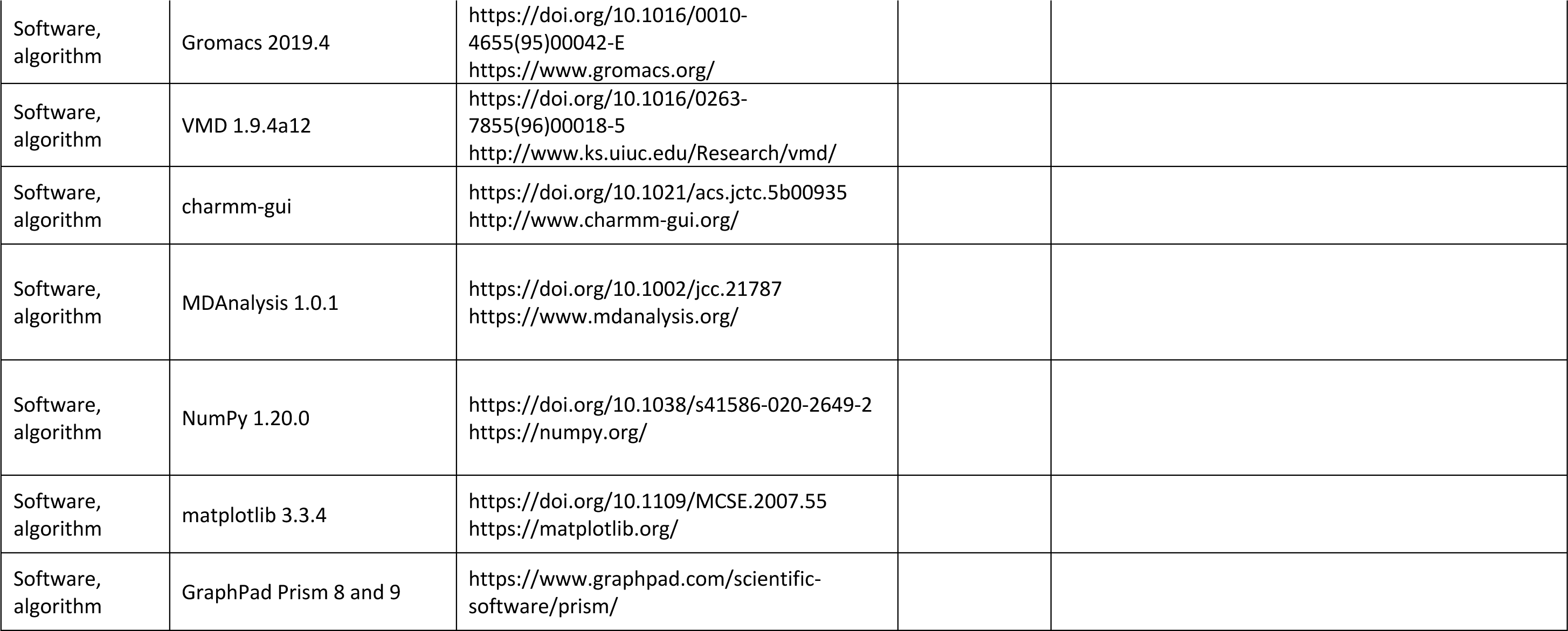
Key resources table.

Each system was minimized using the steepest descents method, then equilibrated with positional restraints on heavy atoms for 100 ps in the NPT ensemble at 310 K with the V-rescale thermostat and a 1 ps coupling time constant, and semi-isotropic Parrinello-Rahman pressure coupling at 1 atm, with a 5 ps coupling time constant (Bussi et al., 2007; Parrinello and Rahman, 1981). Production simulations were run using 2 fs time steps over 50 ns, with 3 repeats run for each state. The simulations were kept relatively short to preserve the conformation of the input structures, whilst allowing sufficient conformational flexibility to sample the side chain motions and rearrangements within the given state. Removal of KdpA and KdpC did not appear to reduce the stability of KdpBF, as all systems had moderate-low backbone RMSDs at the end of the simulations (see Table 3).

Contact analysis was performed by counting the number of residues from each domain which were within 0.4 nm of a residue from a different domain, for every frame over 3 x 50 ns simulation. The domains were defined as the following residues of KdpB: A domain = 89 - 214, N domain = 314 - 450, P-domain = 277 - 313 and 451 - 567. Contact analysis was run with the Gromacs tool gmx select.

High-frequency contacting residue pairs were identified as any pair of residues in contact for at least 90% of frames over 3 x 50 ns of simulation time. Analyses were run using MDAnalysis (Michaud-Agrawal et al., 2011) and plotted using NumPy (Harris et al., 2020) and Matplotlib (Hunter, 2007).

All simulations were run in Gromacs 2019 (Berendsen et al., 1995).

### Figure preparation

All figures were prepared using USCF Chimera (Pettersen et al., 2004), UCSF ChimeraX (Pettersen et al., 2021), VMD (Humphrey et al., 1996), OriginPro 2016, and GraphPad Prism 8 and 9.

## Data availability

Data supporting the findings of this manuscript are available from the corresponding authors upon request. The three-dimensional cryo-EM densities and corresponding modelled coordinates of KdpFABC have been deposited in the Electron Microscopy Data Bank and the Protein Data Bank under the accession numbers summarized in Table 3. The depositions include maps calculated with higher b-factors, both half-maps and the mask used for the final FSC calculation.

## Acknowledgements

CP thanks Michiel Punter for IT support. JMS thanks Paul J.N. Böhm for assistance with cloning and cell growth. PJS acknowledges the University of Warwick Scientific Computing Research Technology Platform for computational access. The electron microscopy within this work is part of the research program National Roadmap for Large-Scale Research Infrastructure (NEMI), project number 184.034.014, which is financed by the Dutch Research Council (NWO).

## Author contributions

Jakob M Silberberg: Validation, Formal analysis, Investigation, writing - original draft, writing – review & editing, visualization

Charlott Stock: Validation, Formal analysis, Investigation, writing - original draft, writing – review & editing, visualization

Lisa Hielkema: Validation, Formal analysis, Investigation, writing - original draft, writing – review & editing, visualization

Robin A Corey: Validation, Formal analysis, Investigation, writing - original draft, writing – review & editing, visualization

Dorith Wunnicke: Validation, Formal analysis, Investigation Jan Rheinberger: Validation, Formal analysis, Investigation Victor RA Dubach: Validation, Formal analysis, Investigation

Phillip J Stansfeld: Conceptualization, Writing – review & editing, Supervision, Project administration, Funding acquisition

Inga Hänelt: Conceptualization, Writing – review & editing, Supervision, Project administration, Funding acquisition

Cristina Paulino: Conceptualization, Writing – review & editing, Supervision, Project administration, Funding acquisition

## Supplementary Data

**Figure 1 – Figure Supplement 1:**
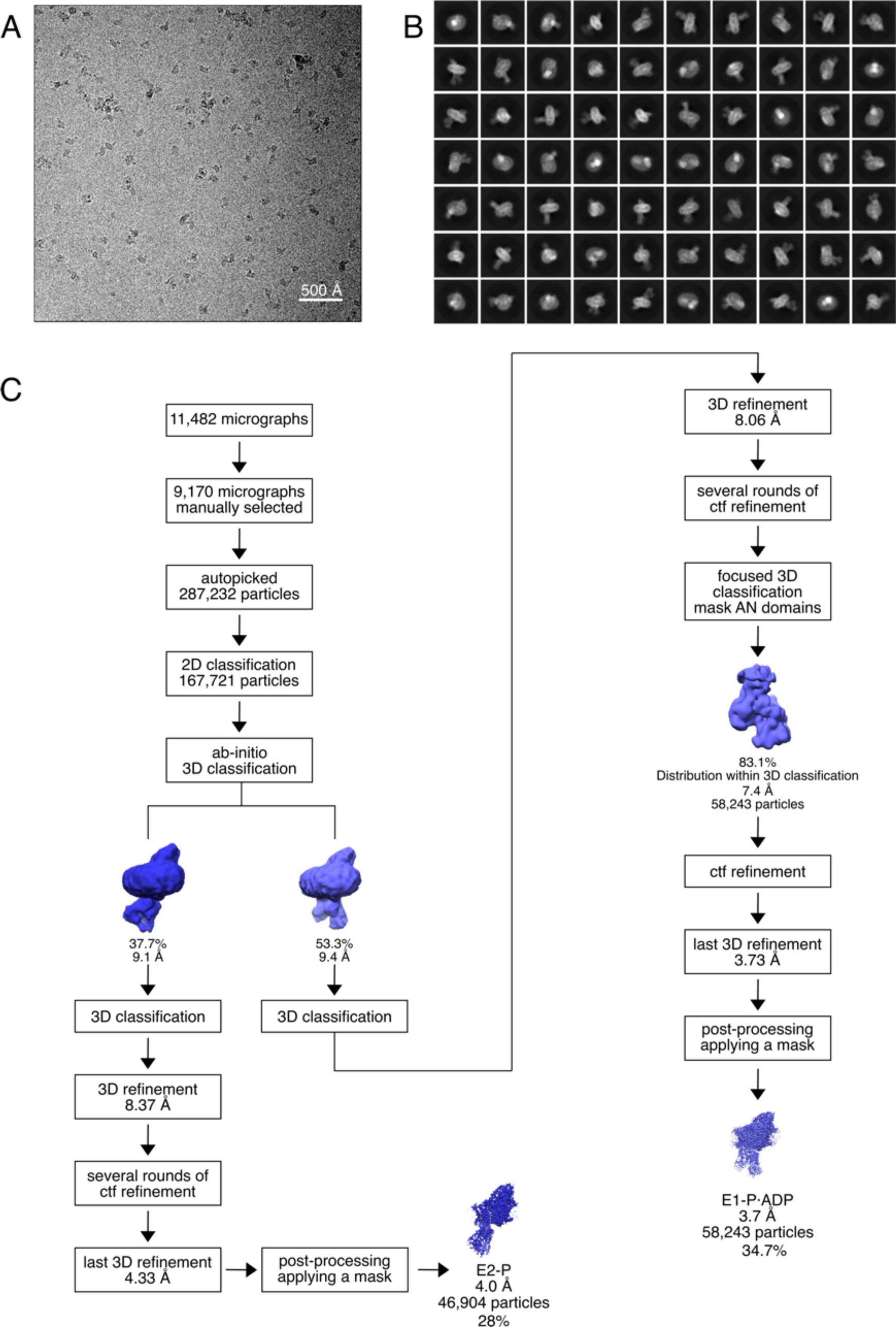
Cryo-EM analysis of the KdpFABS162AC complex under turnover conditions, resulting in E1-P·ADP and E2-P states. **A** Representative cryo-EM image of the recorded data and **B** 2D class averages of vitrified WT KdpFAB_S162-PC_ in the presence of 2 mM ATP and 50 mM KCl. **C** Image processing workflow as described in the material and methods. If not otherwise noted, indicated class percentages refer to the initial set of particles defined after 2D classification.

**Figure 1 – Figure Supplement 2:**
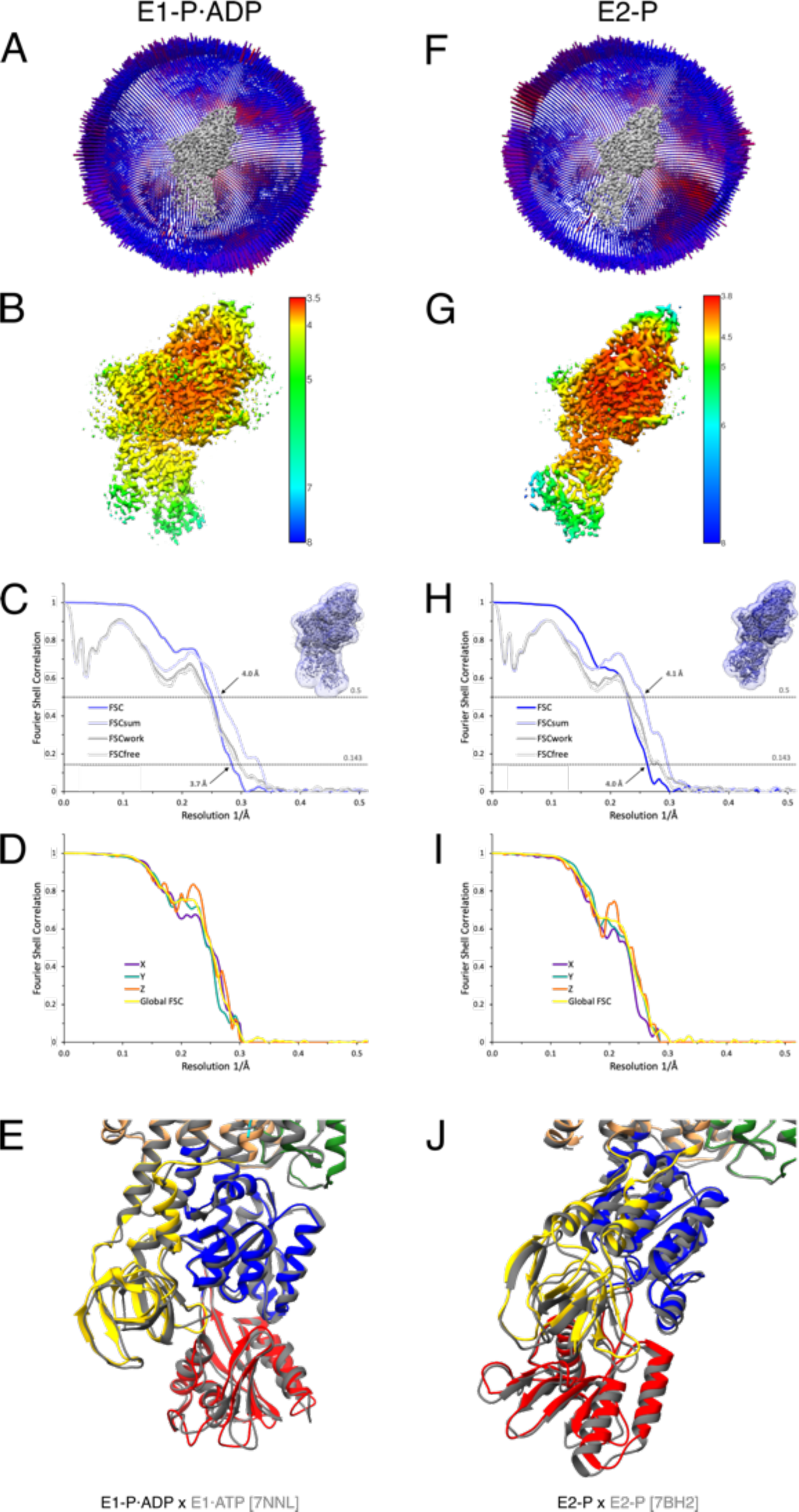
Cryo-EM validation of KdpFABS162AC under turnover conditions. Validation shown for the E1-P·ADP (**A**-**E**) and the E2-P state (**F**-**J**). **A**,**F** Angular distribution plots of particles included in the final unsymmetrized 3D reconstruction. The number of particles with the respective orientations is represented by length and color of the cylinders (long and red: high number of particles; short and blue: low number of particles). **B**,**G** Final reconstruction maps colored by local resolution as estimated by RELION. **C**,**H** FSC plots used for resolution estimation and model validation. The gold-standard FSC plot between two separately refined half-maps is shown in dark blue and indicates final resolutions of 3.7 Å and 4.0 Å for the E1-P·ADP state (**C**) and the E2-P state (**H**), respectively. The FSC model validation curves for FSC_sum_, FSC_work_ and FSC_free_ are described in material and methods and show no overfitting. Thumbnails of the mask used for FSC calculation overlaid on the maps are shown in the upper right corner of both curves. Dashed lines indicate the FSC thresholds used for FSC (0.143) and for FSC_sum_ (0.5). **D**,**I** Anisotropy estimation plots of the final maps show no significant anisotropy. **E**,**J** Superposition of the E1-P·ADP conformation (colored) with the AMPPCP-stabilized E1·ATP structure [7NNL] (gray), and the E2-P conformation (colored) with the BeF_3_^-^-stabilized E2-P structure [7BH2] (gray), respectively, verifying the assignment of each structure at its position in the conformational cycle.

**Figure 1 – Figure Supplement 3:**
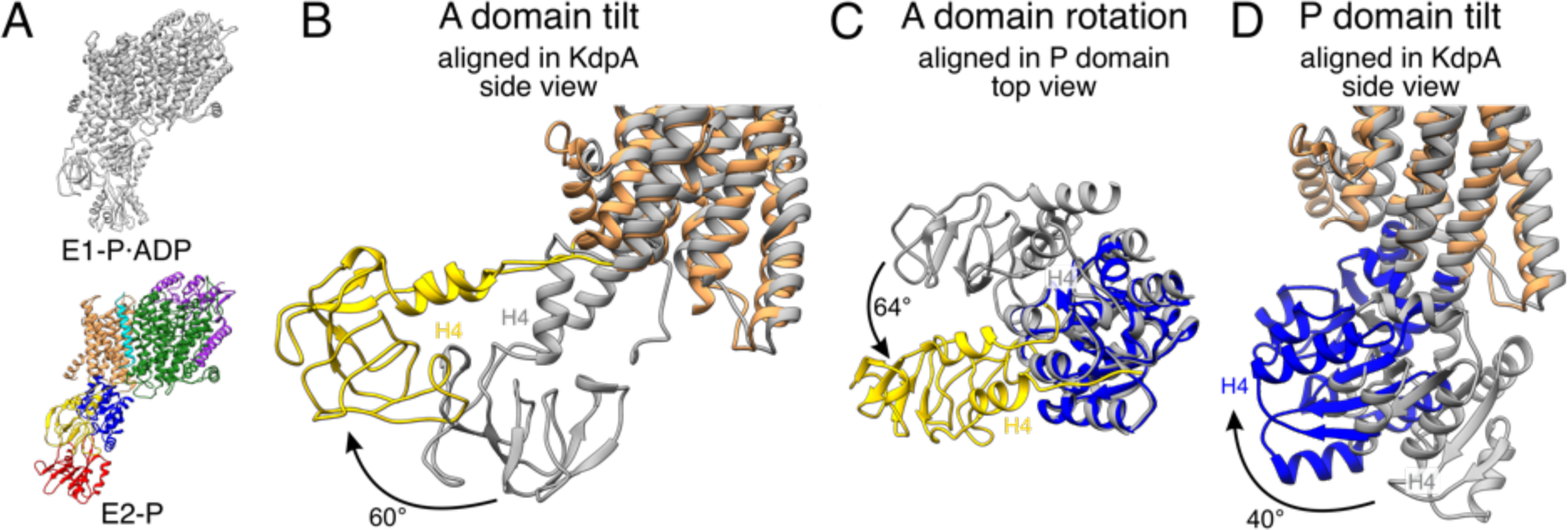
A and P domain movements during the E1-P/E2-P transition. **A** KdpFABS162AC structures in the E1-P·ADP (colors) and E2-P (gray) states. Individual panels show (**B**) the tilt of the A domain (structures aligned on the static KdpA, N and P domain removed for clarity), (**C**) the rotation of the A domain around the P domain (structures aligned on the P domain, N domain and the TMD removed for clarity), and (**D**) the tilt of the P domain (structures aligned on the static KdpA, A and N domain removed for clarity). In the E1-P/E2-P transition, the A domain tilts by 60°, while the P domain tilts by 40°. Additionally, the A domain rotates by 64° around the P domain. Helices from the A and P domains used for angle measurements are labelled in all panels.

**Figure 2 – Figure Supplement 1:**
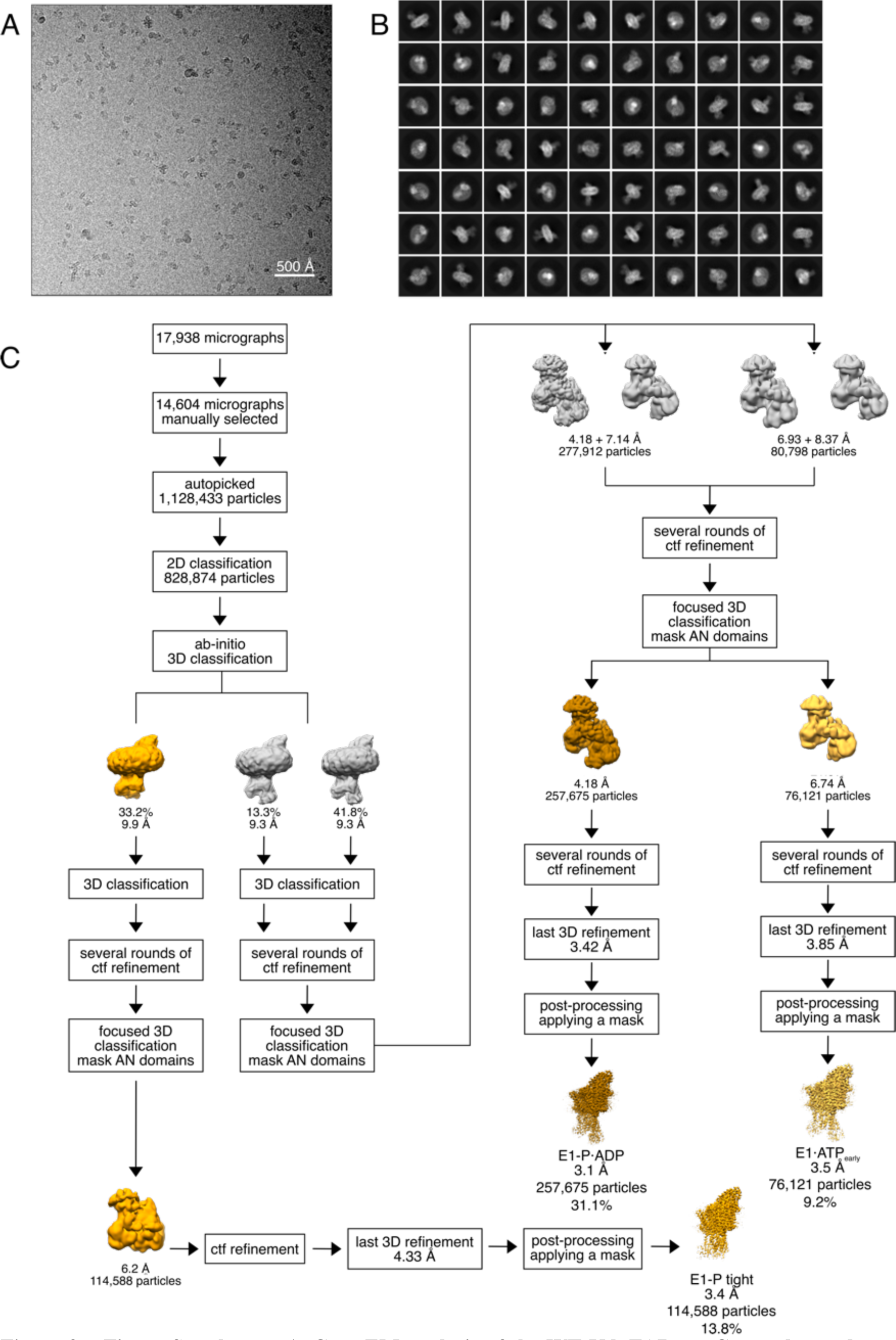
Cryo-EM analysis of the WT KdpFAB_S162-PC_ complex under turnover conditions, resulting in the E1-P tight, E1-P·ADP and E1·ATP_early_ states. **A** Representative cryo-EM image of the recorded data and **B** 2D class averages of vitrified WT KdpFAB_S162-PC_ in the presence of 2 mM ATP and 50 mM KCl. **C** Image processing workflow as described in the material and methods. Indicated class percentages refer to the initial set of particles defined after 2D classification.

**Figure 2 – Figure Supplement 2:**
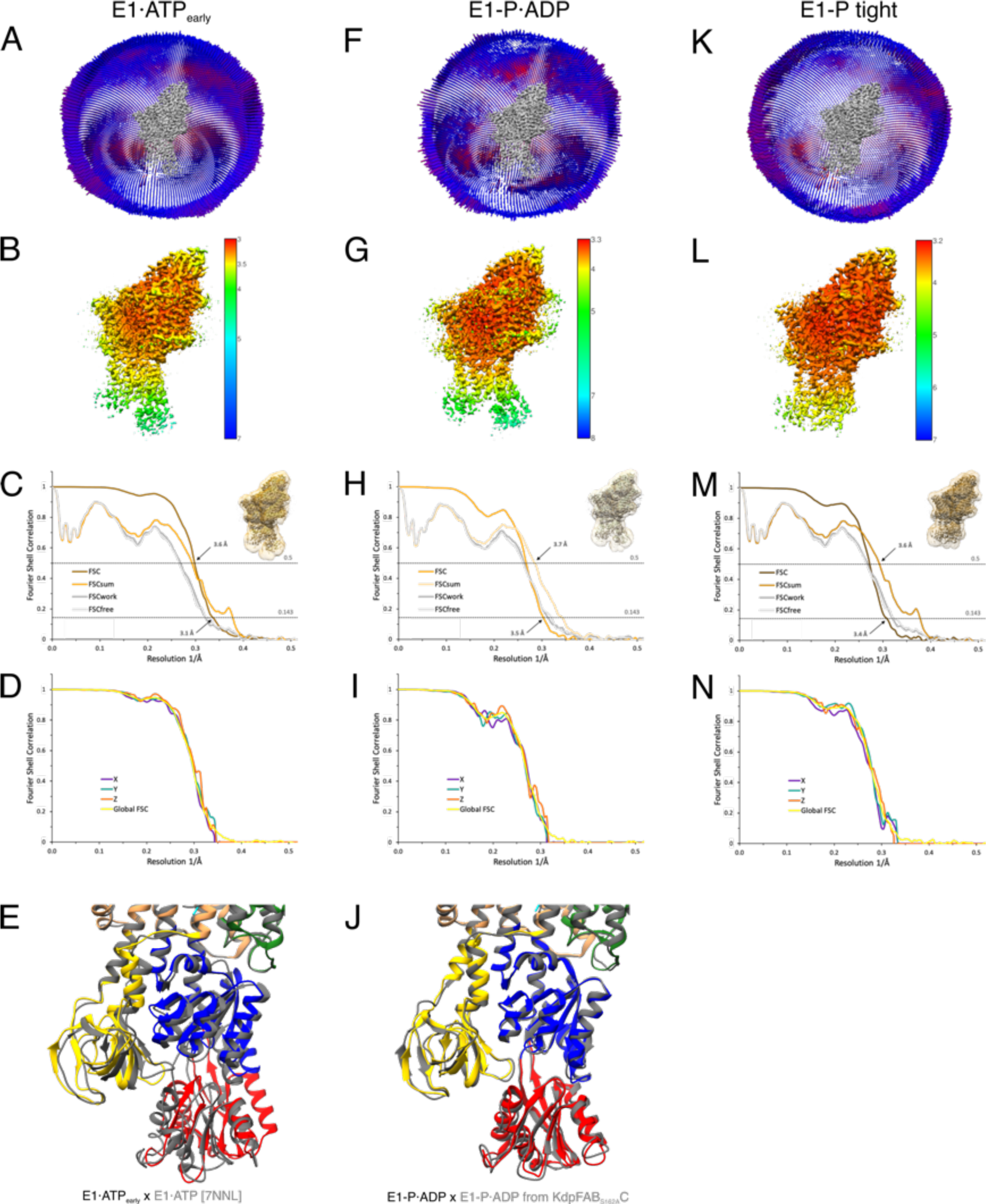
Cryo-EM validation of WT KdpFAB_S162-PC_ under turnover. Validation shown for the E1·ATP_early_ state (**A-E**), the E1-P·ADP state (**F-J**), and the E1-P tight state (**K**-**N**). **A**,**F**,**K** Angular distribution plots of particles included in the final unsymmetrized 3D reconstruction. The number of particles with the respective orientations is represented by length and color of the cylinders (long and red: high number of particles; short and blue: low number of particles). **B**,**G**,**L** Final reconstruction maps colored by local resolution as estimated by RELION. **C**,**H**,**M** FSC plots used for resolution estimation and model validation. The gold-standard FSC plot between two separately refined half-maps indicates final resolutions of 3.5 Å for the E1·ATP_early_ state (C), 3.1 Å for the E1-P·ADP state (**H**) and 3.4 Å for the E1-P tight state (**M**). The FSC model validation curves for FSC_sum_, FSC_work_ and FSC_free_ are described in material and methods and show no overfitting. Thumbnails of the masks used for FSC calculation overlaid on the maps are shown in the upper right corner of the curves. Dashed lines indicate the FSC thresholds used for FSC (0.143) and for FSC_sum_ (0.5). **D**,**I**,**N** Anisotropy estimation plots of the final maps show no significant anisotropy. **E** Superposition of the E1·ATP_early_ structure (colored) with the AMPPCP-stabilized E1·ATP structure [7NNL] (gray), showing a slightly more open N domain in the structure obtained under turnover conditions. **J** Superposition of the E1-P·ADP structure (colored) with the same conformation obtained from KdpFABS162AC under turnover conditions (gray), showing the conformational identity.

**Figure 2 – Figure Supplement 3:**
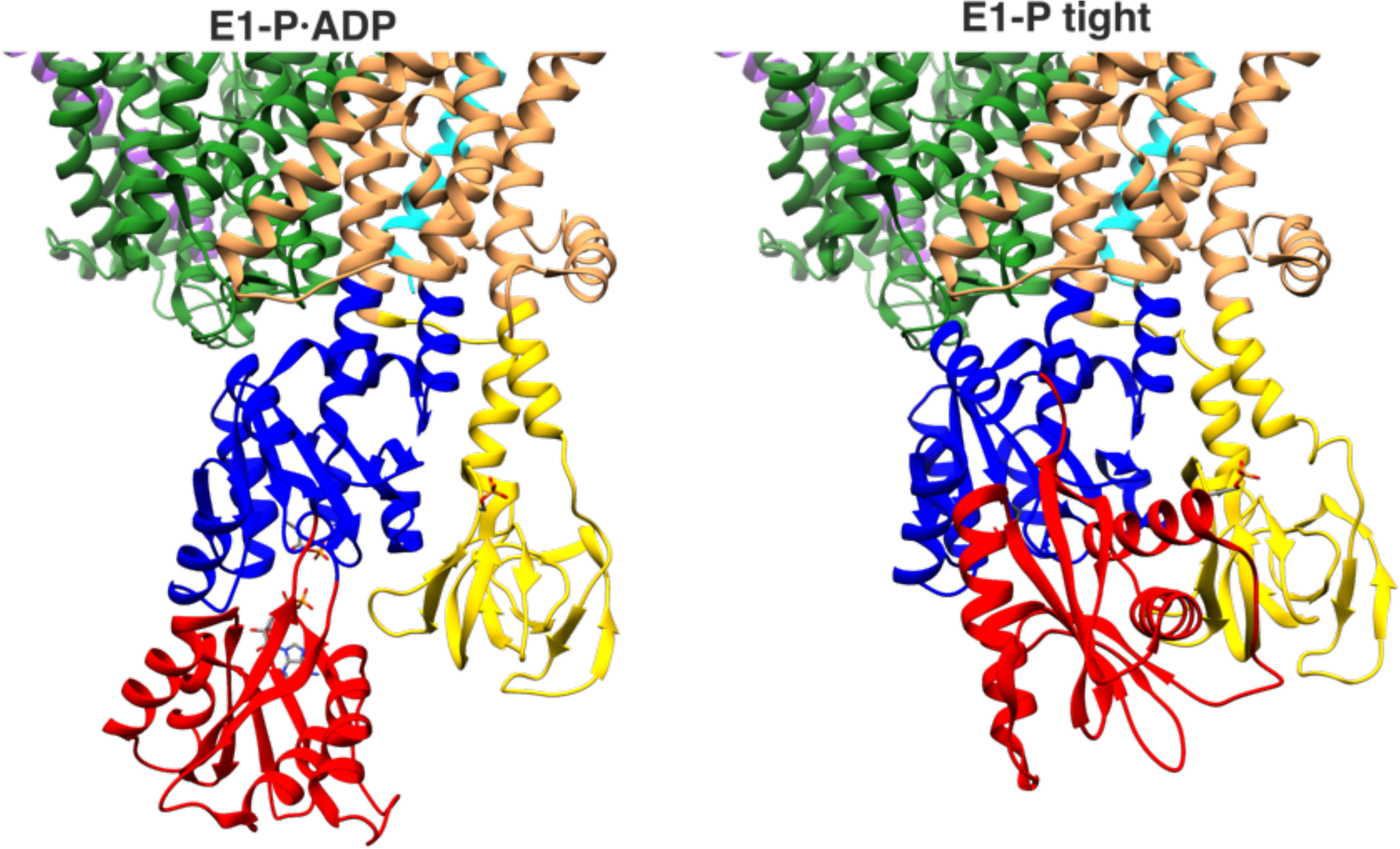
Proximity of A and N domains in E1-P·ADP and E1-P tight KdpFABC. In the E1-P tight state, the interface of the N and A domains is significantly increased compared to the low proximity observed in the E1-P·ADP state.

**Figure 2 – Figure Supplement 4:**
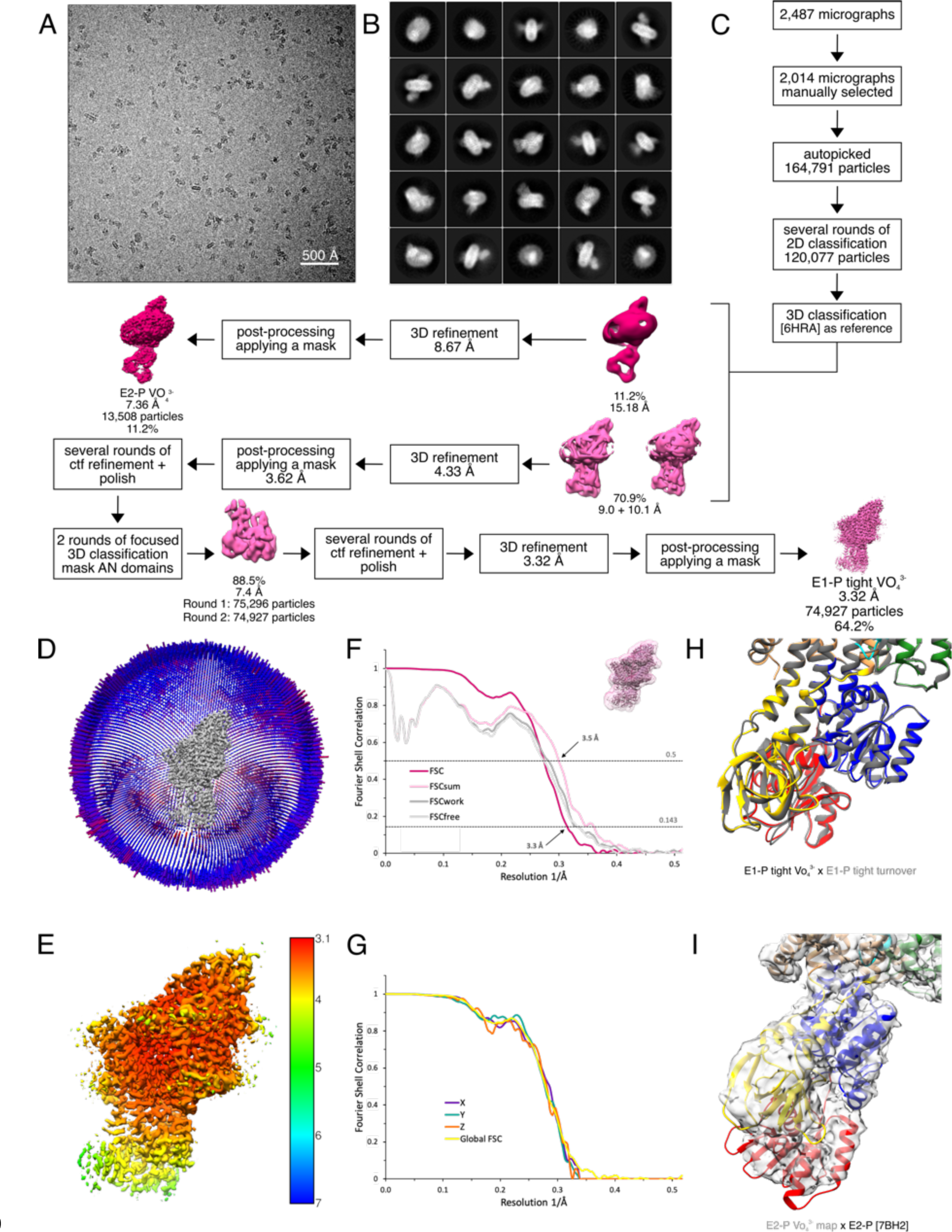
Cryo-EM analysis of WT KdpFAB_S162-PC_ in the presence of orthovanadate, resulting in the E1-P tight and E2-P state. **A** Representative cryo-EM image of the recorded data. **B** 2D class averages of vitrified WT KdpFAB_S162-PC_ in the presence of orthovanadate. **C** Image processing workflow as described in the material and methods section. Indicated class percentages refer to the initial set of particles defined after 2D classification. **D** Angular distribution plot of particles included in the unsymmetrized 3D reconstruction for KdpFABC. The number of particles with the respective orientation is represented by length and color of the cylinders (long and red: high number of particles; short and blue: low number of particles). **E** Final reconstruction map of the E1-P tight state colored by local resolution as estimated by RELION. **F** FSC plot used for resolution estimation and model validation of the E1-P tight state. The gold-standard FSC plot between two separately refined half-maps is shown in red and indicates a final resolution of 3.3 Å. The FSC model validation curves for FSC_sum_, FSC_work_ and FSC_free_ are described in the methods and show no overfitting. A thumbnail of the mask used for FSC calculation overlaid on the map is shown in the upper right corner. Dashed lines indicate the FSC thresholds used for FSC (0.143) and for FSC_sum_ (0.5). **G** Anisotropy estimation plot of the final E1-P tight map, showing no significant anisotropy. **H** Superposition of the E1-P tight structure obtained in the presence of orthovanadate (colored) with the E1-P tight structure of KdpFAB_S162-PC_ obtained under turnover conditions (gray), showing that the adopted conformation is identical. **I** Fit of the BeF_3_^-^-stabilized E2-P structure [7BH2] (colored) into the E2-P map obtained in the presence of orthovanadate (gray), verifying the assignment of the E2-P conformation.

**Figure 2 – Figure Supplement 5:**
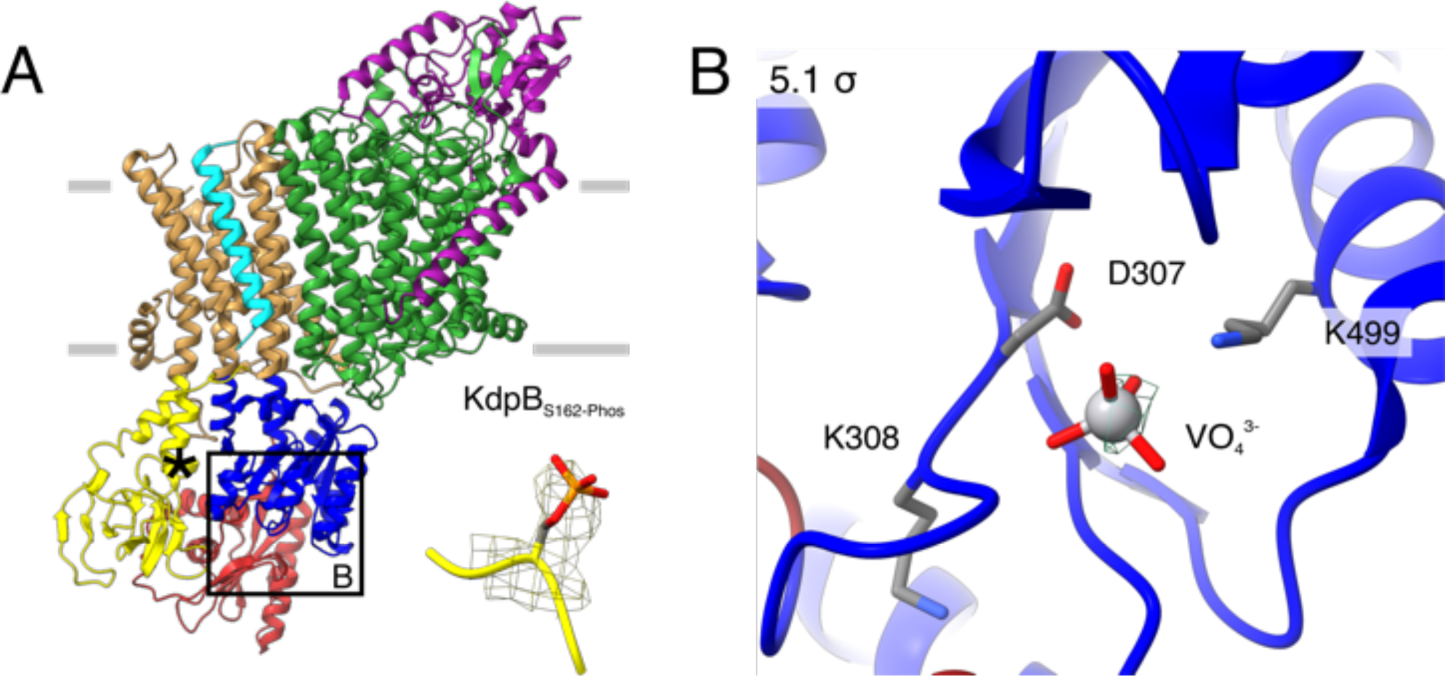
E1-P tight structure of WT KdpFAB_S162-PC_ in the presence of orthovanadate. **A** The E1-P tight structure obtained from KdpFAB_S162-PC_ in the presence of orthovanadate with the KdpB_S162_ phosphorylation highlighted. **C** Nucleotide binding site in the P domain of the E1-P tight conformation, showing the coordinated orthovanadate (VO4^3-^) that mimics phosphorylation of the catalytic KdpB_D307_. Densities are shown at the indicated σ level.

**Figure 3 – Figure Supplement 1:**
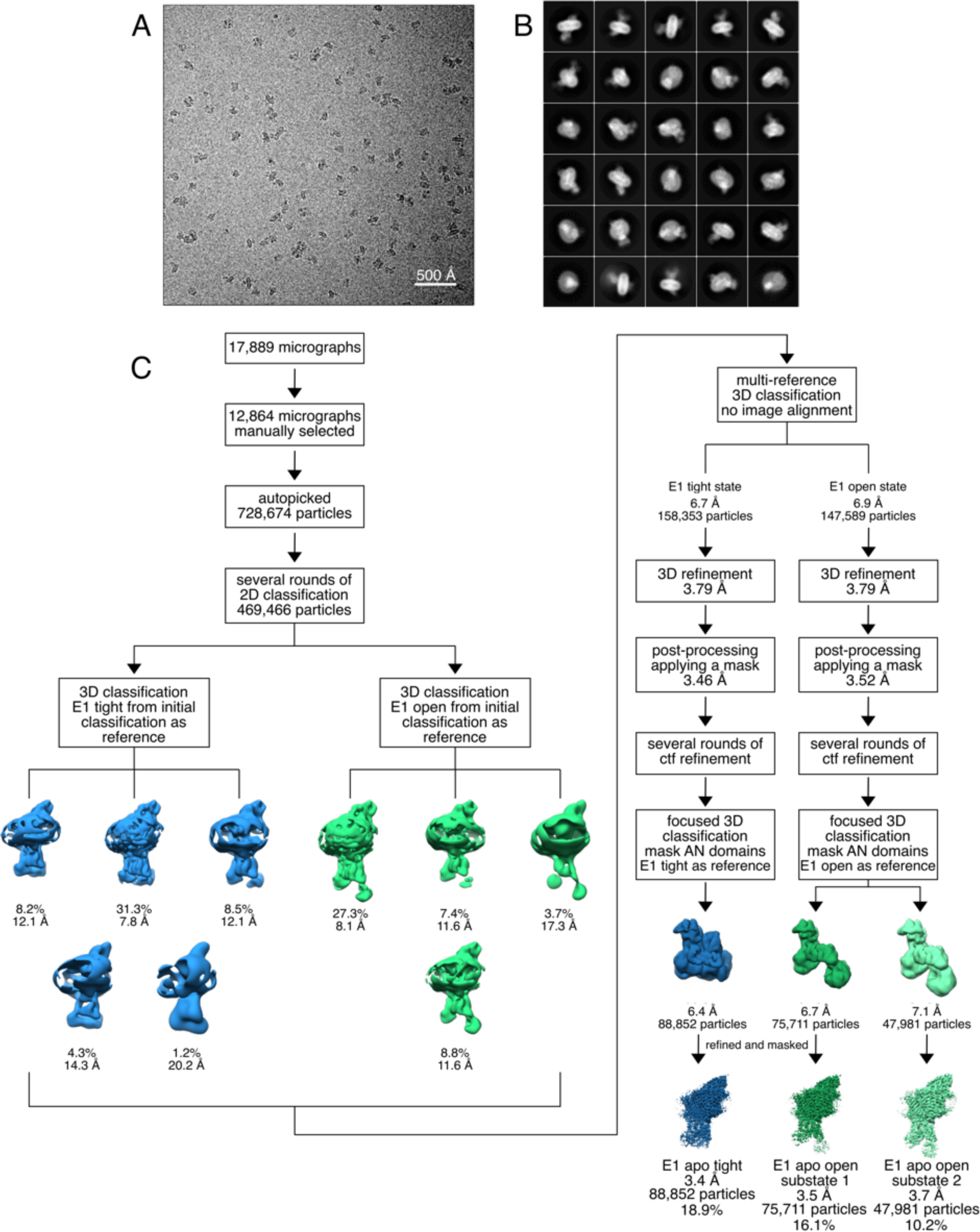
Cryo-EM analysis of the KdpFABS162-P/D307NC complex in the absence of nucleotide, resulting in the E1 apo tight and E1 apo open states. **A** Representative cryo-EM image of the recorded data. **B** 2D class averages of vitrified KdpFABS162-P**/**D307NC in the presence of 50 mM KCl. **C** Image processing workflow as described in the material and methods. Indicated class percentages refer to the initial set of particles defined after 2D classification.

**Figure 3 – Figure Supplement 2:**
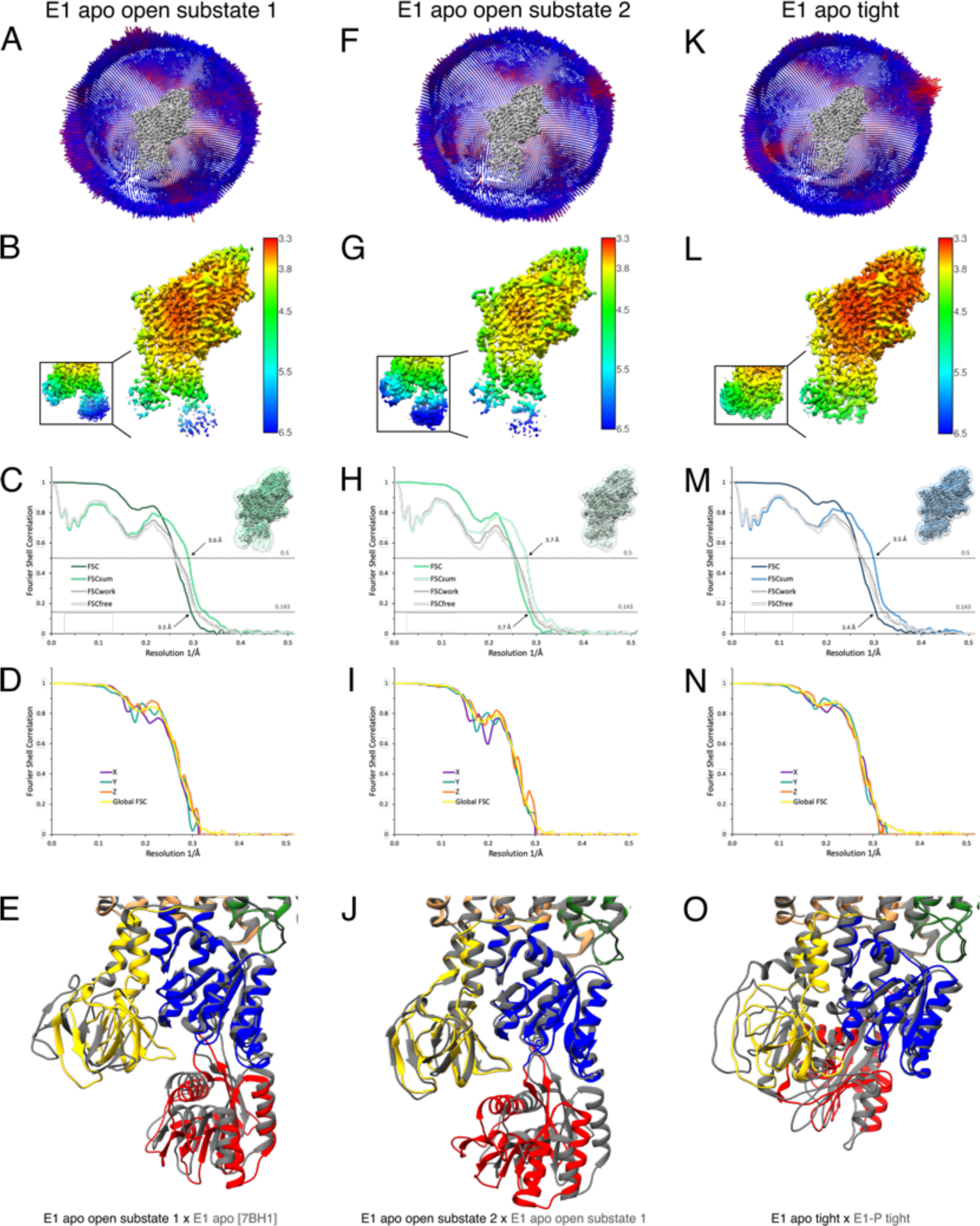
Cryo-EM validation of WT KdpFABS162-P/D307NC under nucleotide-free conditions. Validation shown for E1 apo open substate 1 (**A**-**E**), E1 apo open substate 2 (**F**-**J**) and the E1 apo tight state (**K**-**O**). **A**,**F,K** Angular distribution plot of particles included in the final unsymmetrized 3D reconstruction. The number of particles with the respective orientations is represented by length and color of the cylinders (long and red: high number of particles; short and blue: low number of particles). **B**,**G**,**L** Final reconstruction maps colored by local resolution as estimated by RELION. Inset shows N, P, and A domains at higher contour level. **C**,**H,M** FSC plots used for resolution estimation and model validation. The gold-standard FSC plot between two separately refined half-maps indicates final resolutions of 3.5 Å for the E1 apo open substate 1 (**C**), 3.7 Å for the E1 apo open substate 2 (**H**), and 3.4 Å for the E1 apo tight state (**M**). The FSC model validation curves for FSC_sum_, FSC_work_ and FSC_free_ are described in material and methods and show no overfitting. Thumbnails of the masks used for FSC calculation overlaid on the map are shown in the upper right corner of the curves. Dashed lines indicate the FSC thresholds used for FSC (0.143) and for FSC_sum_ (0.5). **D**,**I**,**N** Anisotropy estimation plots of the final maps show no significant anisotropy. **E** Superposition of E1 apo open substate 1 (colored) with the E1 apo structure [7BH1] (gray), verifying the conformational assignment. **J** Superposition of the E1 apo open substate 2 (colored) with E1 apo open substate 1 (gray), showing a slightly different arrangement of the N domain. **O** Superposition of the E1 apo tight state with the E1-P tight state obtained under turnover conditions, showing a different arrangement of the cytosolic domains.

**Figure 4 – Figure Supplement 1:**
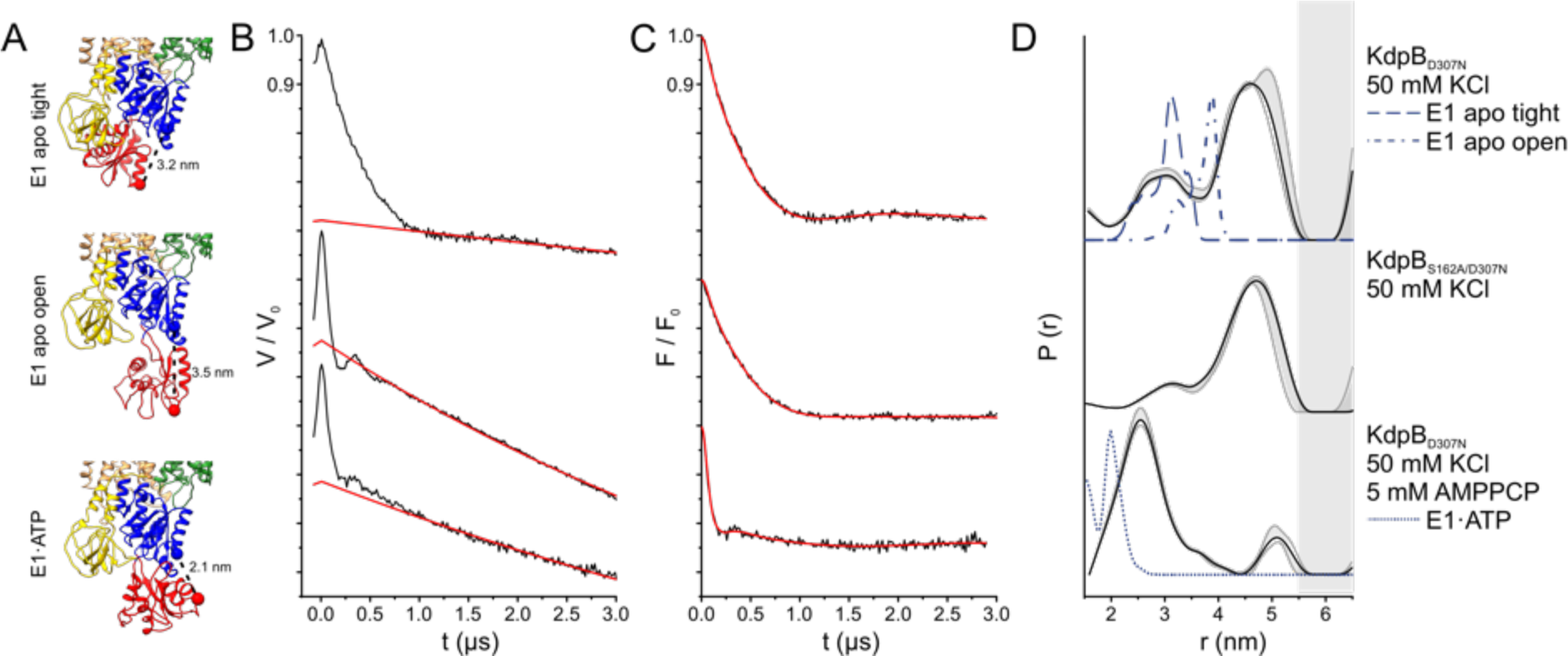
Pulsed EPR measurements of KdpFABC variants in the absence and presence of AMPPCP. **A** Cytoplasmic domains of the anticipated states with expected Cα-Cα distances indicated. **B** Experimental raw data V(t) with fitted background function (red). **C** Background-corrected dipolar evolution function F(t) with applied fit (red). **D** Interspin distance distribution P(r) (black curves) obtained by Tikhonov regularization. Gray background curves indicate the full variation of possible distance distributions. The lower and upper error estimates (blue lines) represent the respective mean value minus and plus two times its standard deviation, respectively; the larger the deviations, the less reliable the predicted distances. Dashed gray lines represent the predicted distance distributions of the E1 apo tight, the E1 apo open and the E1·ATP states, respectively, using the MMM rotamer library analysis. Gray shaded areas starting at 5.5 nm indicate unreliable distances.

**Figure 4 – Figure Supplement 2:**
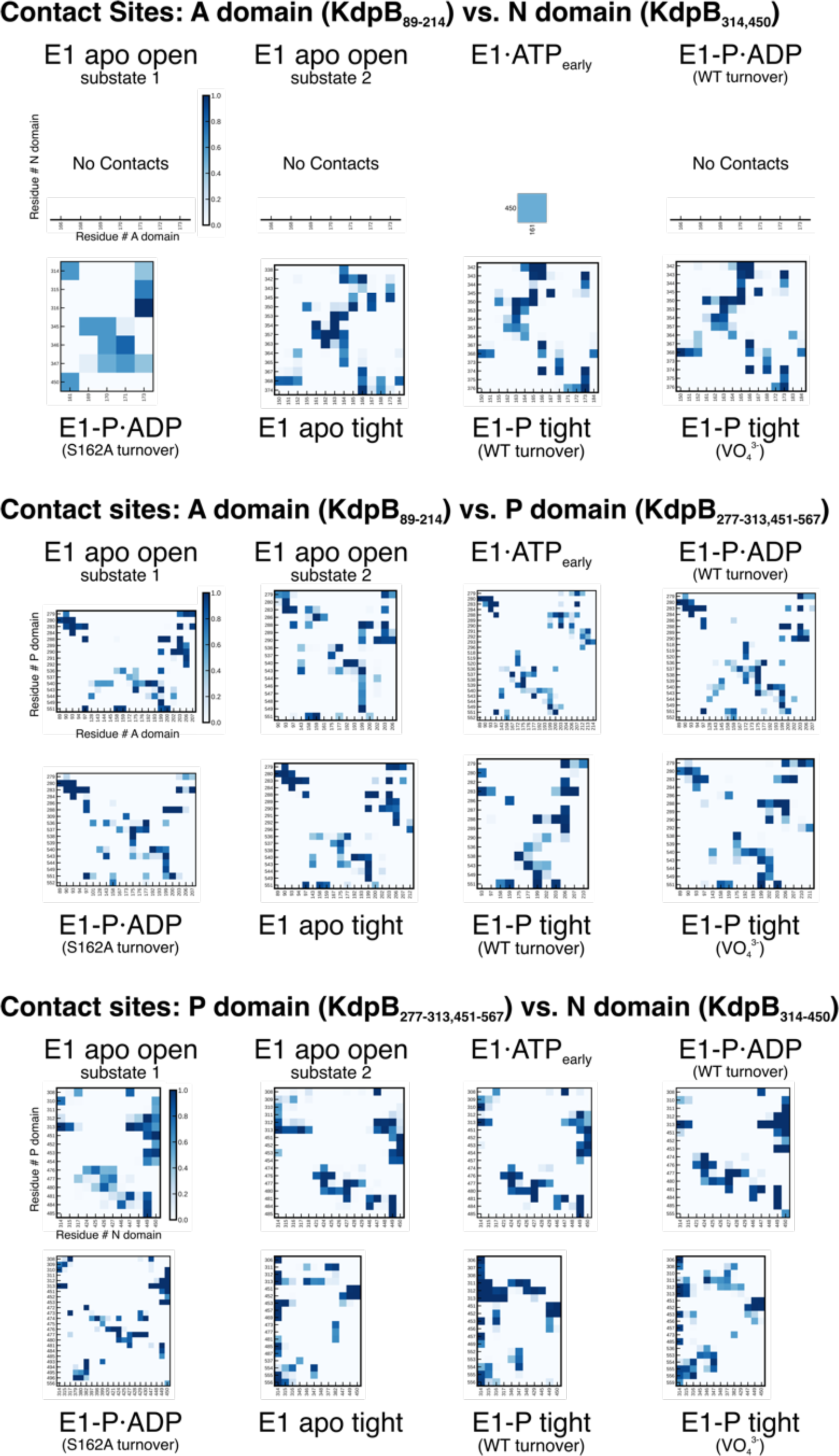
KdpB_S162-P_ mediates stabilization of the E1-P tight state. **A** Contact site analysis of the KdpB N, P and A domains in the E1 conformations of KdpFABC. For each structure, graphs show contact sites observed over 3x50 ns MD simulations, where darker squares represent longer contact in the simulation time. Empty x-axes indicate no contacts.

**Figure 5 – Figure Supplement 1:**
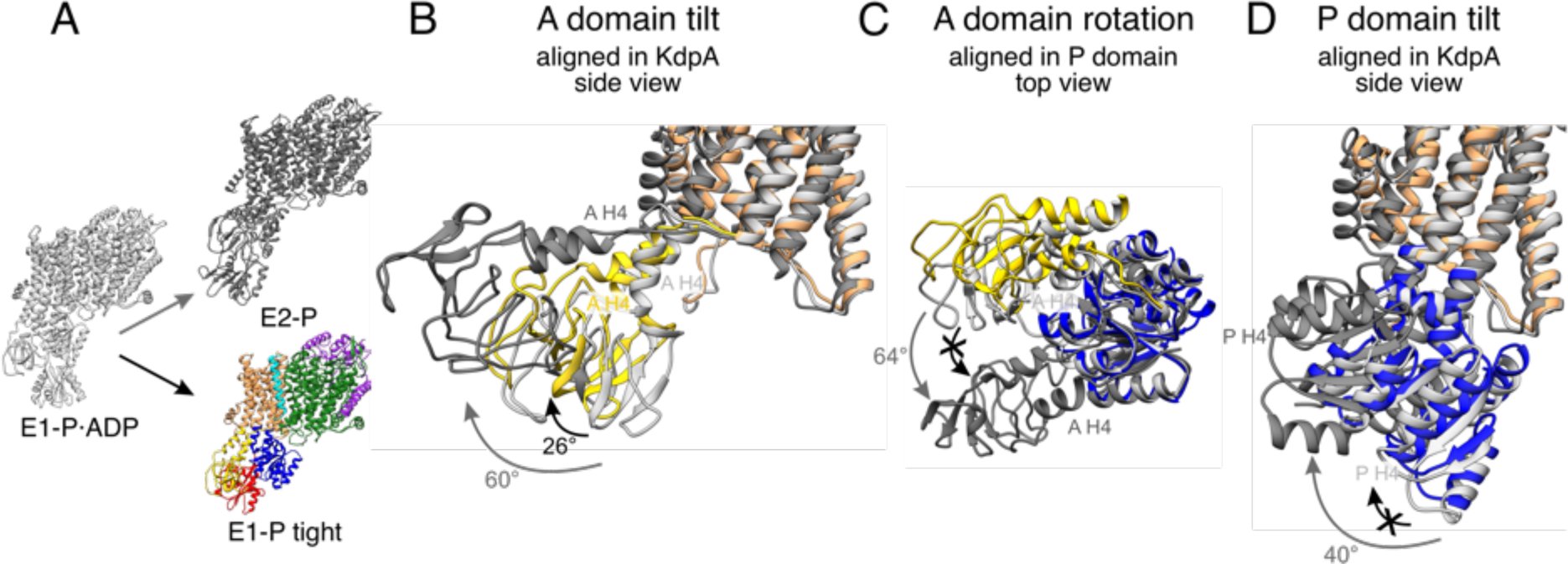
Stalling of A and P domain movements of the E1-P/E2-P transition in the E1-P tight state. **A** Comparison of KdpFAB_S162-PC_ in the E1-P tight state (colored) with KdpFABS162AC in the E1-P·ADP (light gray) and E2-P (dark gray) states. Individual panels show (**B**) the tilt of the A domain (structures aligned on the static KdpA, N and P domain removed for clarity), (**C**) the rotation of the A domain around the P domain (structure of the cytosolic domains aligned on the P domain, N domain and the TMD removed for clarity), and (**D**) the tilt of the P domain (structures aligned on the static KdpA, A and N domain removed for clarity). During the E1-P/E2-P transition (gray arrows), the A domain tilts by 60°, while the P domain tilts by 40°. Additionally, the A domain rotates by 64° around the P domain. The E1-P tight state shows a partial ‘attempted’ transition (black arrows), with a tilt of the A domain by 26°, although the P domain does not undergo global rearrangements and the A domain does not complete its rotation around the P domain. Helices from the A and P domains used for angle measurements are labelled in all panels.

**Figure 5 – Figure Supplement 2:**
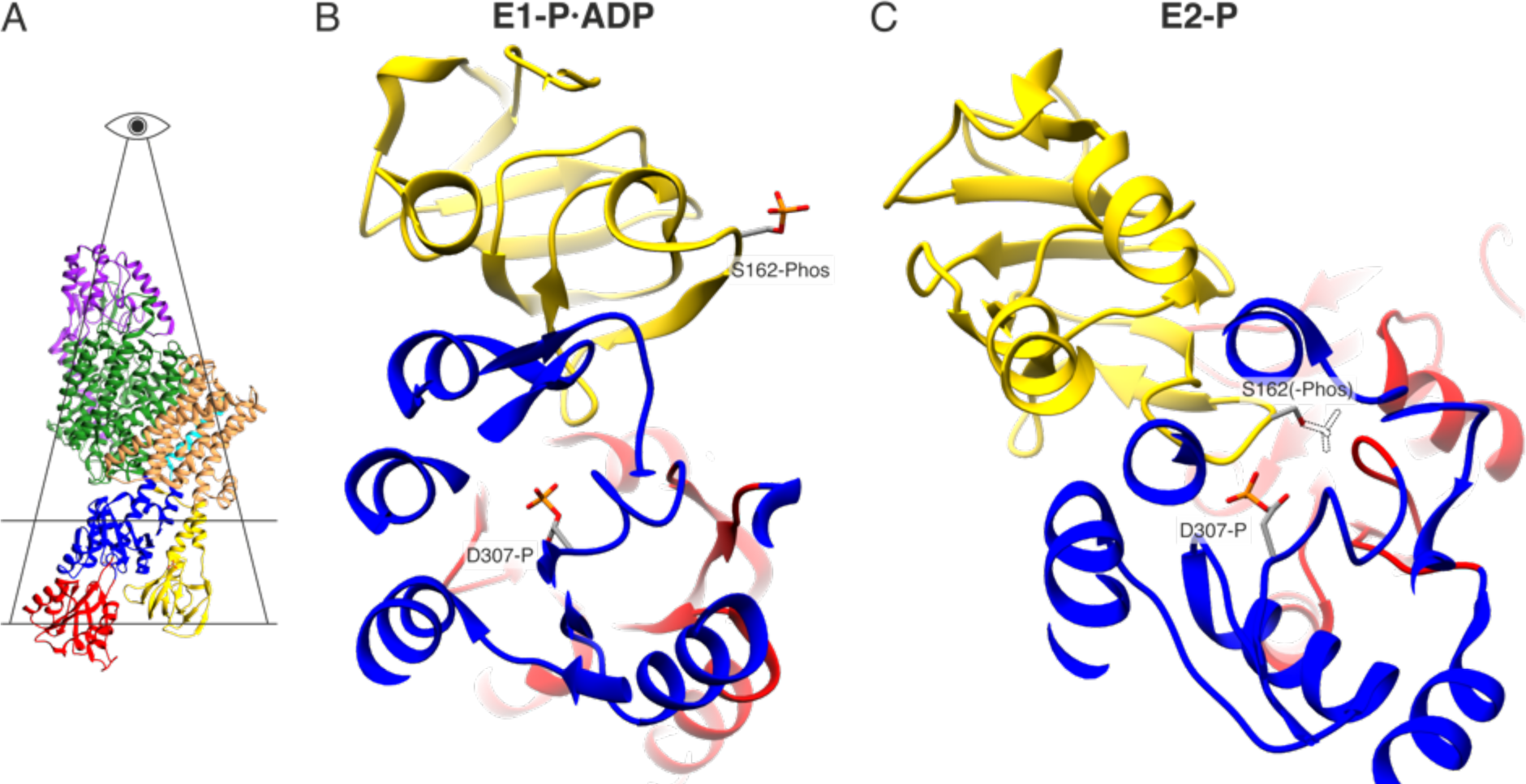
Electrostatic repulsion of KdpB_S162-P_ and KdpB_D307-P_ during the E1-P/E2-P transition. **A** Highlighted section displayed in **B**. **B** View of the proximity between KdpB_D307-P_ and KdpB_S162-P_ in the E1-P·ADP and hypothetical E2-P states. The transition to the E2-P state is not possible in the inhibited KdpFAB_S162-PC_, since the catalytic phosphate would come in close proximity with the inhibitory KdpB_S162-P_, whose theoretical position in the E2-P state is shown as a dashed outline.

**Table 1 – Figure Supplement 1:**
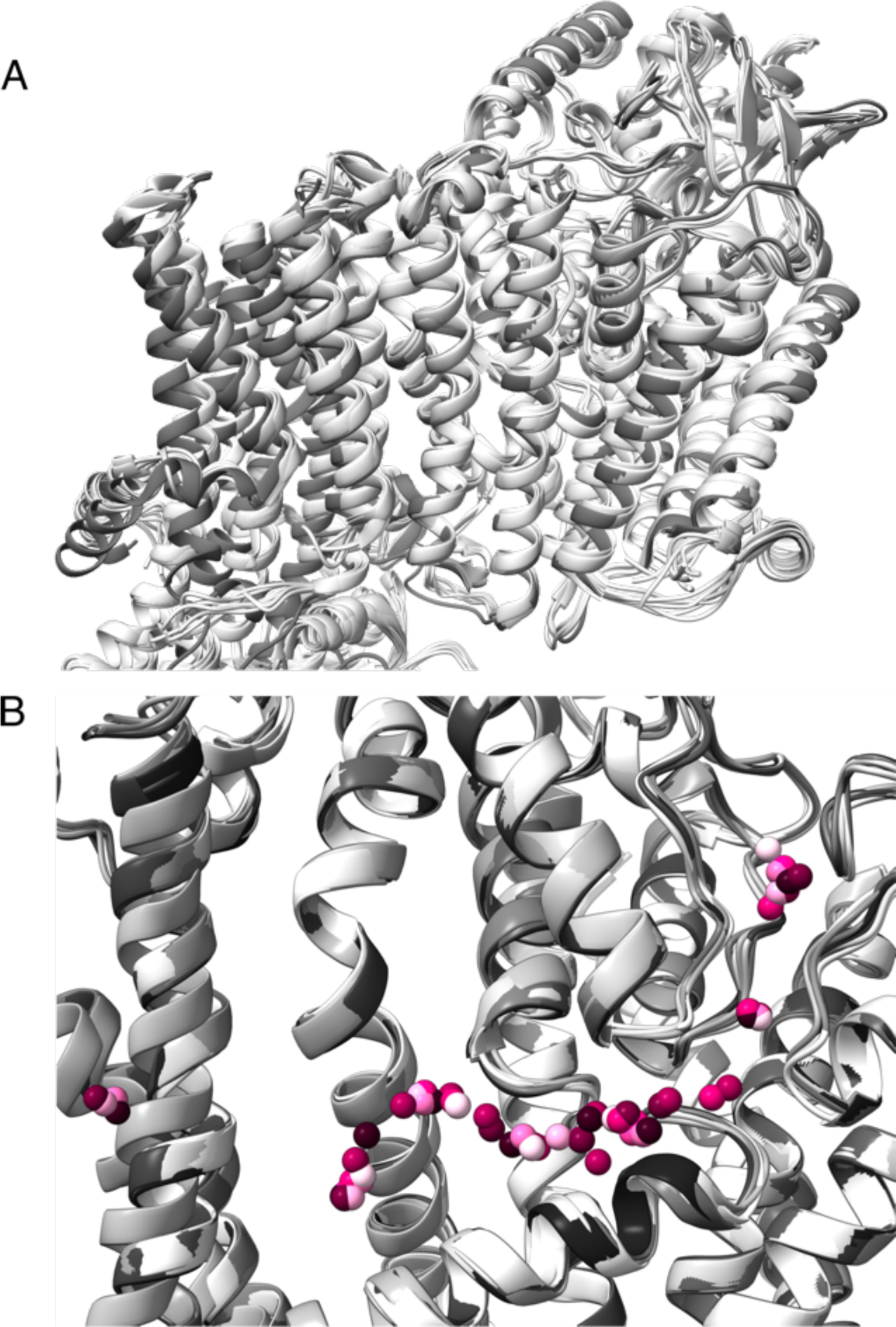
**Comparison of TMDs of all KdpFABC structures obtained in this study. A** Superimposition of all KdpFABC structures obtained in this study on KdpA, showing few significant deviations in the TMD or KdpC. All E1 conformations shaded in light gray, all E2 conformations in dark gray. **B** Superimposition of all potential potassium ions in the intersubunit tunnel found in all structures presented in this paper. Ions for each structure are presented in a different pink shade.

**Table 1 – Table Supplement 1:** Cryo-EM data collection, refinement, and validation statistics

|  | KdpFABC<br>E1-ATP<sub>early</sub> | KdpFABC<br>E1-P-ADP | KdpFABC<br>E1-P tight | KdpFABC<br>E1-P tight<br>(VO<sub>4</sub><sup>3-</sup>) | KdpFABC<br>E2-P<br>(VO<sub>4</sub><sup>3-</sup>) | KdpFAB<sub>S162A</sub>C<br>E1-P-ADP | KdpFAB<sub>S162A</sub>C<br>E2-P | KdpFAB<sub>D307N</sub>C<br>E1 apo tight | KdpFAB<sub>D307N</sub>C<br>E1 apo open 1 | KdpFAB<sub>D307N</sub>C<br>E1 apo open 2 |
| --- | --- | --- | --- | --- | --- | --- | --- | --- | --- | --- |
|  | EMD-14913<br>[7ZRG] | EMD-14917<br>[7ZRK] | EMD-14912<br>[7ZRE] | EMD-14911<br>[7ZRD] | EMD-14347 | EMD-14919<br>[7ZRM] | EMD-14918<br>[7ZRL] | EMD-14914<br>[7ZRH] | EMD-14915<br>[7ZRI] | EMD-14916<br>[7ZRJ] |
| **Data collection and processing** |  |  |  |  |  |  |  |  |  |  |
| Magnification | 49,407 | 49,407 | 49,407 | 49,407 | 49,407 | 49,407 | 49,407 | 49,407 | 49,407 | 49,407 |
| Voltage (keV) | 200 | 200 | 200 | 200 | 200 | 200 | 200 | 200 | 200 | 200 |
| Electron exposure (e<sup>-</sup>/Å<sup>2</sup>) | 52 | 52 | 52 | 52 | 52 | 52 | 52 | 52 | 52 | 52 |
| Defocus range (μm) | -0.5 to -2.0 | -0.5 to -2.0 | -0.5 to -2.0 | -0.5 to -2.0 | -0.5 to -2.0 | -0.5 to -2.0 | -0.5 to -2.0 | -0.5 to -2.0 | -0.5 to -2.0 | -0.5 to -2.0 |
| Pixel size (Å) | 1.012 | 1.012 | 1.012 | 1.012 | 1.012 | 1.012 | 1.012 | 1.012 | 1.012 | 1.012 |
| Symmetry imposed | C1 | C1 | C1 | C1 | C1 | C1 | C1 | C1 | C1 | C1 |
| Initial particle images (no.) | 1,128,433 | 1,128,433 | 1,128,433 | 164,891 | 164,891 | 287,232 | 287,232 | 728,674 | 728,674 | 728,674 |
| Final particle images (no.) | 76,121 | 257,675 | 114,488 | 74,927 | 13,508 | 58,243 | 46,904 | 88,852 | 75,711 | 47,981 |
| Map resolution (Å) | 3.5 | 3.1 | 3.4 | 3.3 | 7.4 | 3.7 | 4.0 | 3.4 | 3.5 | 3.7 |
| FSC threshold | 0.143 | 0.143 | 0.143 | 0.143 | 0.143 | 0.143 | 0.143 | 0.143 | 0.143 | 0.143 |
| Map resolution range (Å) | 3.3-6.5 | 3.0-5.5 | 3.1-5.0 | 3.1-5.0 | N/A | 3.5-6.5 | 3.8-7.0 | 3.3-5.5 | 3.4-6.5 | 3.6-6.5 |
| **Refinement** |  |  |  |  |  |  |  |  |  |  |
| Initial model used (PDB code) | 7NNL | 7NNL | 6HRA | 6HRA | N/A | 7NNL | 6HRB | 6HRA | 6HRA | 6HRA |
| Model resolution (Å) | 3.7 | 3.6 | 3.6 | 3.5 | N/A | 4.0 | 4.1 | 3.5 | 3.6 | 3.7 |
| FSC threshold | 0.5 | 0.5 | 0.5 | 0.5 | N/A | 0.5 | 0.5 | 0.5 | 0.5 | 0.5 |
| Model resolution range (Å) | 80-3.5 | 80-3.1 | 80-3.4 | 80-3.3 | N/A | 80-3.7 | 80-4.0 | 80-3.4 | 80-3.5 | 80-3.7 |
| Sharpening B-factor (Å<sup>2</sup>) | -132 | -122 | -134 | -55 | -195 | -123 | -160 | -113 | -117 | -119 |
| Model composition |  |  |  |  |  |  |  |  |  |  |
| Non-hydrogen atoms | 11097 | 11103 | 11071 | 11072 | N/A | 10890 | 10782 | 11065 | 11065 | 11065 |
| Protein residues | 1456 | 1456 | 1456 | 1456 | N/A | 1456 | 1456 | 1456 | 1456 | 1456 |
| Ligands | K: 7<br>CDL: 2<br>ATP: 1 | K: 13<br>CDL: 2<br>ADP: 1<br>MG: 1 | K: 9<br>CDL: 2 | K: 9<br>CDL: 2<br>VO4: 1 | N/A | K: 5<br>ADP: 1<br>MG: 1 | K: 1 | K: 7<br>CDL: 2 | K: 7<br>CDL: 2 | K: 7<br>CDL: 2 |
| B factors (Å<sup>2</sup>) |  |  |  |  |  |  |  |  |  |  |
| Protein | 55.0 | 54.3 | 50.2 | 23.8 | N/A | 54.8 | 69.9 | 33.0 | 52.5 | 47.5 |
| Ligand | 54.8 | 55.6 | 40.1 | 25.9 | N/A | 116.6 | 64.4 | 29.6 | 49.5 | 37.8 |
| R.m.s. deviations |  |  |  |  |  |  |  |  |  |  |
| Bond lengths (Å) | 0.006 | 0.006 | 0.007 | 0.006 | N/A | 0.004 | 0.005 | 0.004 | 0.004 | 0.005 |
| Bond angles (°) | 0.849 | 0.877 | 0.976 | 0.893 | N/A | 0.838 | 0.908 | 0.828 | 0.820 | 0.908 |
| Validation |  |  |  |  |  |  |  |  |  |  |
| MolProbity Score | 1.74 | 1.72 | 1.70 | 1.53 | N/A | 1.68 | 1.77 | 1.73 | 1.68 | 1.80 |
| Clash score | 7.24 | 8.04 | 7.79 | 5.88 | N/A | 6.82 | 8.17 | 7.43 | 6.90 | 9.34 |
| Poor rotamers, % | 0.00 | 0.00 | 0.00 | 0.00 | N/A | 0.00 | 0.00 | 0.00 | 0.00 | 0.00 |
| Ramachandran plot |  |  |  |  |  |  |  |  |  |  |
| Favored (%) | 95.02 | 95.91 | 95.98 | 96.68 | N/A | 95.64 | 95.26 | 94.43 | 95.71 | 95.64 |
| Allowed (%) | 4.98 | 4.09 | 4.02 | 3.32 | N/A | 4.36 | 4.74 | 4.57 | 4.29 | 4.36 |
| Outliers (%) | 0.00 | 0.00 | 0.00 | 0.00 | N/A | 0.00 | 0.00 | 0.00 | 0.00 | 0.00 |

